# Quantum machine learning for untangling the real-world problem of cancers classification based on gene expressions

**DOI:** 10.1101/2023.08.09.552597

**Authors:** Mohadeseh Zarei Ghoabdi, Elaheh Afsaneh

## Abstract

Quantum machine learning algorithms using the power of quantum computing provide fast-developing approaches for solving complicated problems and speeding-up calculations for big data. As such, they could effectively operate better than the classical algorithms. Herein, we demonstrate for the first time the classification of eleven cancers based on the gene expression values with 4495 samples using quantum machine learning. In addition, we compare the obtained quantum classification results with the classical outcomes. By implementing a dimensional reduction method, we introduce significant biomarkers for each cancer. In this research, we express that some of the identified gene biomarkers are consistent with DNA promotor methylation, and some other ones can be applied for the survival determination of patients.

## 1. Introduction

In recent years, quantum machine learning (QML) has attracted a great deal of attention due to its great potential in improving algorithms for increasing accuracy, speeding up, and using lesser computational resources. QML is a rapidly growing research field at the intersection of quantum computing and machine learning (ML).

Machine learning technologies aim to find patterns and create models through the accomplishment of learning tasks in various areas from scientific research to technical/industrial fields (1, 2) Particularly, machine learning algorithms have been widely used for classifying biological data, identifying novel biomarkers, exploring new disease subtypes, etc. (3, 4). These approaches result in the recognition of functional factors accurately after employing a biological-based feature selection (5–8).

Executing ML for the analysis of big data needs extensive computational operations. To improve the computing process, quantum methods have found great interest for their significant properties in saving time and computational resources. Quantum computations perform calculating operations more efficiently than classical ones based on quantum mechanical principles including superposition, measurement, and entanglement.

In classification tasks, a popular QML algorithm is the Quantum Support Vector Machine (QSVM) which is a quantum version of the classical SVM algorithm. QSVM utilizes quantum principles in computational operations and provides considerable results than the classical SVM. The SVM algorithm separates data points into classes by drawing decision boundaries between them with respect to the maximum margin of hyperplanes. Although QSVM employs similar procedures, it uses quantum kernels to do computations that are hard to perform by classical approaches. In one research, it was discussed that performing the algorithm of quantum-enhanced least-square SVM provides exponential speed-up (9). It was suggested that the difficult classical machine learning (CML) tasks could solve by classically intractable quantum feature maps. In another research, the performance of kernel-based QSVM and quantum neural networks on small practical datasets were analyzed by different quantum feature maps (10). Their proposed QSVM algorithms showed better accuracy than the classical SVM by performing on both a quantum simulator and a real quantum computer. Further in (11), a kernel-based support vector machine was implemented on a DW2000Q quantum annealer. The results showed that the introduced quantum variant of SVM was a proper practical alternative to the standard SVM. On the other hand, through the current noisy intermediate-scale quantum computers (NISQ) era, there are considerable challenges in the advancement of quantum technologies. In this case, one of the main issues is achieving computations with faster speeds than classical computers. Quantum supremacy has been introduced as a significant milestone that could perform superior computations on quantum systems beyond the power of classical ones (12).

Despite the capability of quantum computational supremacy, achieving this potential is still challenging that is because of early quantum computer developments (13). The current quantum devices are extensively under the effect of noise and error connection. In addition, quantum computers are still constructed with a few hundred qubits that are far from quantum supremacy with thousands of qubits. Alongside quantum hardware improvement to reach quantum supremacy, the simulation of quantum algorithms on classical computers has performed significant development (14).

From another point of view, even with the significant improvement of quantum hardware and software, the most quantum advantages performed on the solution of non-real-world problems. It means that the considered challenges mostly have not been focused on real-world practical relevance. One of the major fields for implementing QML algorithms with critical roles in analyzing real-world big data is biological data which includes gene/protein expression data and protein structure (15), healthcare and life sciences (16), and also drug design and drug discovery (17).

In this research, to demonstrate and achieve quantum benefits in investigating real-world problems, we employed a quantum machine learning algorithm for classifying different cancers. Using our suggested approach, we analyzed and classify eleven cancers from healthy samples and could find possible biomarkers. Moreover, we compared the obtained results between the quantum ML algorithms and the classical ML techniques. This comparison showed the good capability of QML in the classification of transcriptomic data. Referring to the limitation of quantum computers with the number of qubits, we performed the quantum machine learning computations using simulation on the classical hardware.

## 2. Methods

In this section, we define our employed quantum machine learning method for classifying eleven various cancer data. We first describe some basic concepts of quantum computing. Then, we introduce quantum machine learning and the related elements including quantum kernels and quantum circuits. For implementing quantum algorithms and utilizing their applications, we focus on SVM and QSVM, and finally, we present the used algorithm, Skqulacs-QSVM.

### 2.1 Quantum Computing

In classical computing, information is stored in bits in the form of either 0 or 1. While in quantum computing, the basic unit of information is stored as quantum bits so-called qubits. To express a qubit in mathematical terms, it is considered a vector in a two-dimensional complex Hilbert space with the computational basis |0⟩ and |1> (18). In contrast to a classical bit, a qubit exists in one of three states |0>, |1> or superposition (linear combination) of basis states as c_0_ |0⟩ + c_1_ |1⟩. An important phenomenon of quantum mechanics is quantum entanglement, which originates from the concept of superposition. Quantum entanglement refers to the correlation of two or more quantum states where each of the correlated states can influence the other ones. However, the particles of states could physically separate at a great distance. With the measurement of one particle, the superposition state collapses which affects instantaneously the other ones. These quantum characteristics imply that more information can be stored in the entangled qubit states than in the individual ones. It provides quantum computing with exponential speedup for solving traditional difficult problems and big data as well.

### 2.2 Quantum Machine Learning

Quantum machine learning benefits from the integration of quantum algorithms within classical machine learning methods for improving computational speed-up and for technological facilities in the current NISQ era. QML algorithms as the quantum version of ML mechanisms have been performed in various strategies (2). In quantum-inspired machine learning techniques, the methods of quantum computing are implemented to improve the classical ML algorithms (19). Quantum-enhanced machine learning algorithms refer to executing classical data on quantum computers (20). The potential of quantum computers can be demonstrated by providing the exponential computational space due to utilizing qubits over the classical computers in the Boolean space. In hybrid classical-quantum machine learning algorithms, classical and quantum methods are merged to enhance the performance of the model (21). In this research, we employ a quantum machine learning algorithm that works based on the support vector machine method for classifying big data cancer. For it, we used simulating the quantum algorithm on a classical computer. In other words, we considered our implemented QML algorithm in the hybrid classical-quantum machine learning algorithms category.

### 2.3 Quantum Support Vector Machine

Quantum support vector machine algorithm utilizes the beneficial aspects of both quantum hardware and software to improve the performance of standard SVM algorithms. Support vector machine is a popular supervised machine learning algorithm that is generally used in classification problems. To classify data, a decision boundary is drawn between data points for determining a hyperplane and differentiating two classes. Finding an optimal hyperplane corresponds to maximizing the distance between support vectors and the hyperplane (4). Particularly, support vectors are used for the data points that are nearest to the hyperplane on the edges of the margin. SVM employs the support vectors to detach samples rather than employing the diversities in class means.

In SVM, kernel methods are used for separating data and also improving classification accuracy. It means that kernels provide mapping data into a higher-dimensional feature space for easier data separation (22). Applying quantum kernels for QML algorithms leads to mapping classical data into the quantum states in Hilbert space using quantum feature maps. The infinite possibility of the dimension of the kernel Hilbert space makes the kernel approach powerful (23).

A quantum kernel is a function determining the resemblance between two quantum states in the feature space (24). Employing a kernel provides QML algorithms to classify quantum states according to their similarities. The advantages of performing quantum kernels have been shown in the enhancement of machine learning algorithms for solving problems (25). In QML, encoding the classical data points x from the input dataset *χ* into the quantum states |1ψ (x)⟩ belonging to the higher dimension Hilbert space ℋ is so-called a quantum feature map (26, 27).

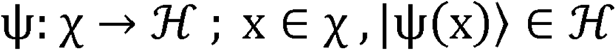

where ψ denotes a quantum feature encoding that acts through a mathematical procedure for mapping data. Quantum kernel counterparts of the classical kernel are implemented depending on kernel function to construct a quantum classifier in QML algorithms. In kernel function κ, the inner product is built between two quantum states representing two mapped data points x, x’ in combination with a quantum feature map defined as:

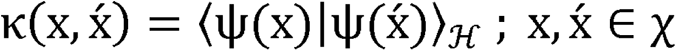

in which, ⟨.|.⟩ denotes the dot product (28). Through applying quantum kernels, the transformation of classical input data into quantum states can be represented in encoding circuit form as (29):

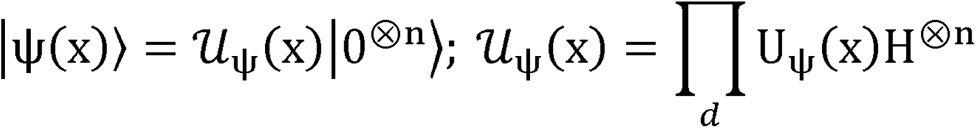

where 𝒰_ψ_(x), a quantum feature encoding circuit is consisting of H as the Hadamard gates and U_ψ_(x) as the unitary operations containing Pauli gates. In which, n shows the number of qubits used for encoding and d represents the depth of the circuit for creating the quantum feature map circuit. In addition, the initial state for n qubits is |0^⊗n^⟩ Il - |10… 0) (Figure 1).

**Figure 1:**
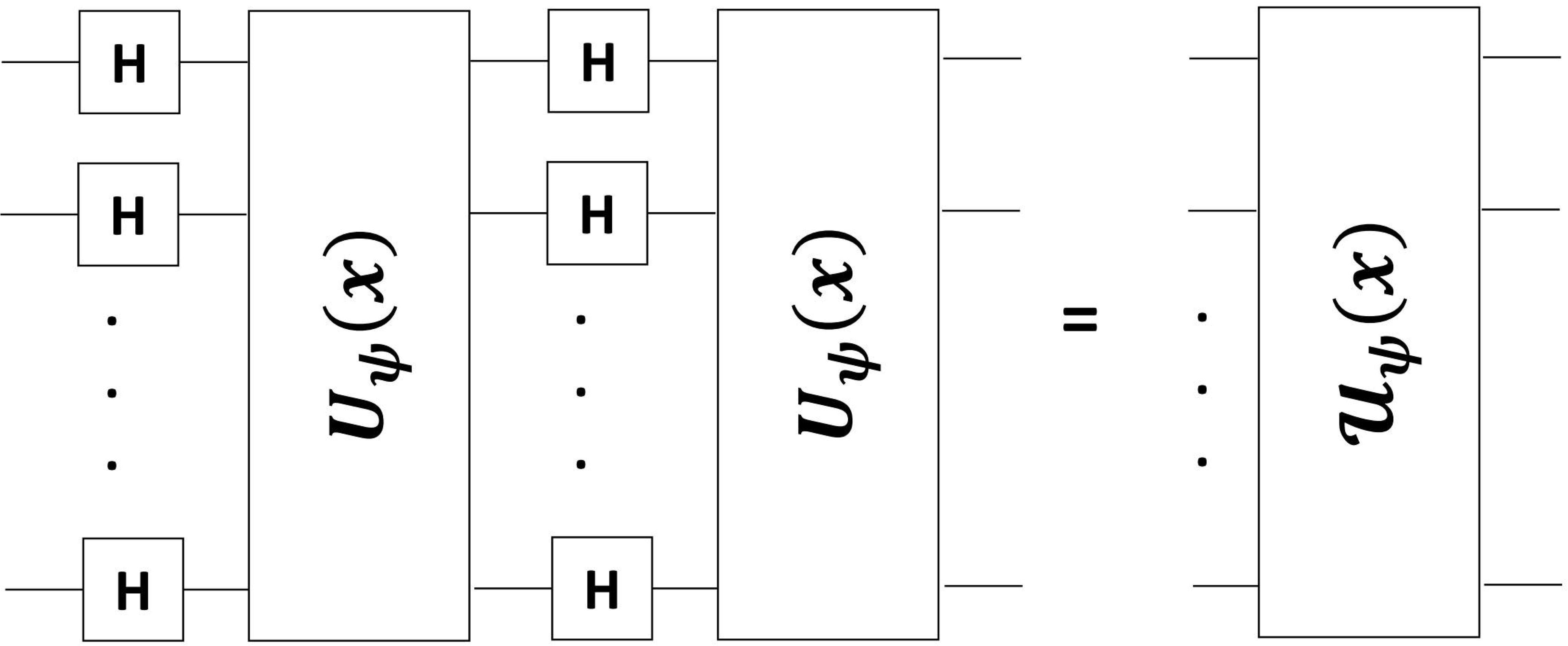
Feature map circuit 𝒰_ψ_(x) consists of Hadamard gate H and unitary operation U_ψ_(x) applying for each single-qubit.

Generally, a quantum circuit in a quantum computing process involves the modification of qubits with the quantum gates that are in combination with the traditional logical operators and gates. Quantum gates as the building blocks of quantum circuits consist of Hadamard gates and controlled Pauli-gates. Hadamard gate brings qubits into a superposition state and controlled Pauli-gates enable qubits to rotate around the x, y, and z-axis for providing quantum entanglement between qubits (10). To estimate an observable value in a quantum circuit, measurement is performed multiple times and the result is obtained as a prediction (30).

### 2.4 Skqulacs-QSVM

In the present research, to classify and analyze different data cancers, we employed the QML scikit-qulacs package (31). This quantum machine learning algorithm originated based on one of the fastest quantum circuit simulators named Qulacs (32). Qulacs is planned for the help of researchers in quantum computing operations since it particularly shows significant speed-up in exploration and performing algorithms with high accuracy. The scikit-qulacs package introduces a quantum machine learning algorithm using the quantum circuit simulator Qulacs. In this QML algorithm, quantum circuits are created using the function “Learning Circuit” from sub-package “skqulacs.circuit” in “skqulacs package” (31). A quantum circuit as an array of quantum gates is represented underlying “ParametricQuantumCircuit” class. In this procedure, quantum gates are generated with the function “create_yzcx_ansatz” (33) from “skqulacs.circuit.pre_defined” sub- package. The provided circuit can be visualized using “circuit_drawer” function in the “qulacsvis” package (34). Through the classification process, standard quantum support vector classification was performed by function “QSVM” from “skqulacs.qsvm” package (35). In this contribution, we implemented “skqulacs.qsvm” package on our data to utilize the advantages of this package in performing classification. We called this quantum support vector machine algorithm as skqulacs-QSVM. In the next sections, we observed the results of applying skqulacs- QSVM algorithm to our prepared data. We carried out this algorithm by simulating quantum circuits with low-depth to take advantage of speeding up.

Furthermore, for skqulacs-QSVM algorithm in the process of feature selection, we used the Fisher score algorithm to select features. This algorithm is a dimensional reduction and feature-ranking approach that has been used for the selection of genes from the expression profile data. The procedure of feature selection is explained as follows. Given X ∈ Rm×n is a matrix of gene expression data (m and n represent the number of genes and samples, respectively) and NG = (U, C, D, δ) is a neighborhood decision system for gene expression data, the Fisher score is calculated by

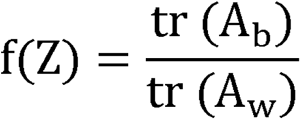

Where tr () denotes the trace of a matrix, A_W_ indicates the scatter matrix within the same class, and A_b_ shows the scatter matrix between the cancer samples and their paired normal samples.

Then, an exploratory strategy is commonly employed to compute the score for each gene using similar criteria. Afterward, the Fisher score of the k-th gene is obtained by:

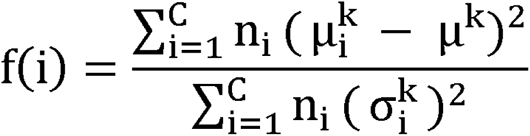

Where n_i_ refers to the sample number of the i-th class. 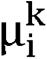 and 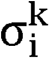 illustrate the mean and standard deviation of the samples from the i-th category with the k-th gene, respectively. Also, μ^J<^ shows the mean magnitude of the k-th gene samples (36, 37). We wrote and executed codes in a Python 3.11.3 environment.

## 3. Data collection

We downloaded GDC TCGA RNA-seq (HTSeq) counts of 11 cancers including Bladder Cancer (BLCA), Colon Adenocarcinoma (COAD), Glioblastoma (GBM), Head and Neck Squamous Cell Carcinoma (HNSC), Kidney Renal Clear Cell Carcinoma (KIRC), Liver hepatocellular carcinoma (LIHC), Lung Adenocarcinoma (LUAD), Prostate Adenocarcinoma (PRAD), Rectal Adenocarcinoma (READ), Stomach Adenocarcinoma (STAD), and Thyroid Cancer (THCA) from UCSC Xena database (https://xenabrowser.net/). A total of 4495 cancer and healthy samples have been employed. The details of the samples are described in Table 1.

**Table 1.**
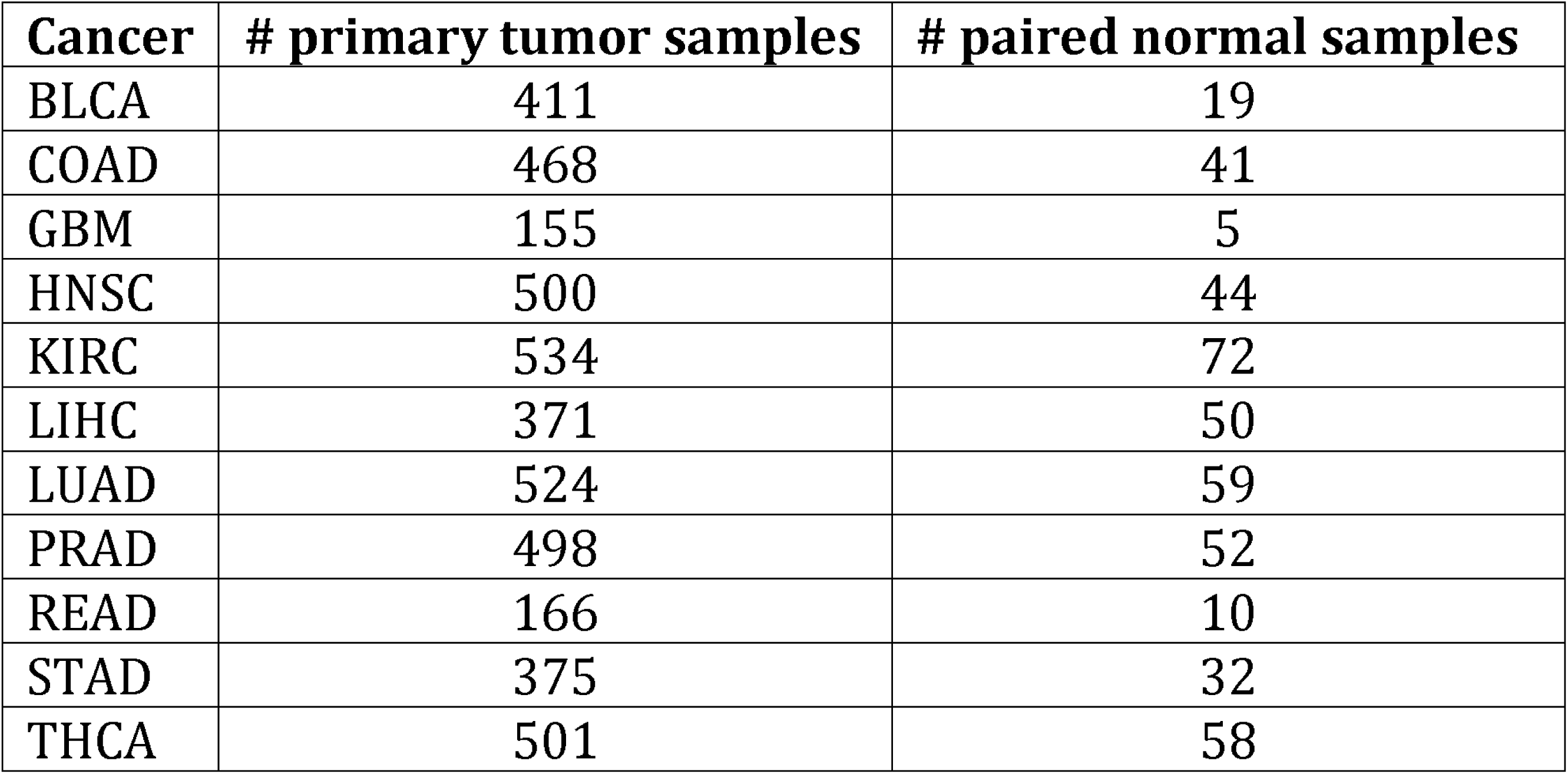
Details of the cancer datasets.

| Cancer | # primary tumor samples | # paired normal samples |
| --- | --- | --- |
| BLCA | 411 | 19 |
| COAD | 468 | 41 |
| GBM | 155 | 5 |
| HNSC | 500 | 44 |
| KIRC | 534 | 72 |
| LIHC | 371 | 50 |
| LUAD | 524 | 59 |
| PRAD | 498 | 52 |
| READ | 166 | 10 |
| STAD | 375 | 32 |
| THCA | 501 | 58 |

The Ensemble IDs were mapped to official gene symbols. The Ensemble IDs without gene names were excluded. The database contains log2 (conut+1) transformed data. For this reason, before starting the gene filtering, we converted them back to integer raw counts.

### 3.1 Gene filtering and determining differentially expressed genes

We removed all rows with zero-sum and rows with sum (cpm(data)>100) < 2. We also discarded genes with variances less than 1st quartile. Afterward, we performed TMM normalization and log2 transformation. To determine differentially expressed genes (DEGs), we applied edgR and limma voom methods using packages edgeR (version 3.36.0) and limma (3.50.0) executed in the R programming environment (version 4.3.1), respectively. We considered the criteria of Benjamini-Hochberg adjusted. P. value < 0.05 (38) and |logFC| >3. Finally, we found the common DEGs identified by the two methods for further analysis. We excluded long-non coding RNAs.

### 3.2 Preparing data for classification

The data were randomly partitioned to the train and test sets in Python 3.11.3 (70/30). To overcome the imbalance between normal and cancer classes in the training dataset, we used Synthetic Minority Over-sampling Technique (SMOTE). SMOTE produces fresh data in feature space by employing the K-Nearest Neighbor (KNN) algorithm (39).

### 3.3 Pathway enrichment analysis

We utilized g:profiler webtools (https://biit.cs.ut.ee/gprofiler) (40) to enrich classifier genes in the Kyoto Encyclopedia of Genes and Genomes (KEGG) and Reactome pathway databases.

## 4. Results

We have implemented classical SVM (CSVM) and skqulacs-QSVM on eleven types of cancer datasets in a classical computer in order to compare their performances. The details of the analysis steps and results are explained in the below sections.

### 4.1 Determination of DEGs

Table 2 indicates the number of genes after several filtering (explained in the Data Collection section) and the number of DEGs identified by edgeR, limma, and common ones between these two methods (Supplementary data file 1). We considered the shared DEGs for further analysis in order to have a strict criterion for the entry of the genes to the machine learning analysis step.

**Table 2.** Number of genes after filtering and number of DEGs.

| **Cancer** | **# Genes** | **# DEGs_edgeR** | **# DEGs_limma** | **# DEGs_common** |
| --- | --- | --- | --- | --- |
| BLCA | 6550 | 337 | 249 | 146 |
| COAD | 5575 | 314 | 305 | 180 |
| GBM | 4612 | 555 | 497 | 438 |
| HNSC | 5792 | 325 | 224 | 154 |
| KIRC | 5761 | 370 | 392 | 239 |
| LIHC | 4842 | 233 | 217 | 64 |
| LUAD | 6675 | 478 | 252 | 151 |
| PRAD | 4876 | 88 | 23 | 12 |
| READ | 4344 | 229 | 231 | 154 |
| STAD | 5858 | 303 | 162 | 86 |
| THCA | 4430 | 177 | 150 | 92 |

### 4.2 Feature selection and performance analysis of CSVM and skqulacs-QSVM

Fisher score is a supervised feature selection approach that chooses features based on their scores. It means that the ranked genes are sorted according to their importance. We used the Fisher score to rank the genes (Supplementary data file 2). To find an optimum number of features that lead to obtaining maximum test accuracy, we tested the first 15 ranked genes for CSVM. To construct our skqulacs-QSVM model, we specified the number of qubits for the first 15 features ranked by Fisher score. The expression amount of each sample was used as input data for both classical and quantum machine learning methods. Figure 2A and 2B shows the obtained test accuracy after employing CSVM and skqulacs-QSVM, respectively, when the different number of ranked features were selected. The results disclosed the optimum number of applied qubits as 3, 3, 4, 12, 8, 9, 11, 12, 4, 8, and 13 for BLCA, COAD, GBM, HNSC, KIRC, LIHC, LUAD, PRAD, READ, STAD, and THCA, respectively.

**Figure 2:**
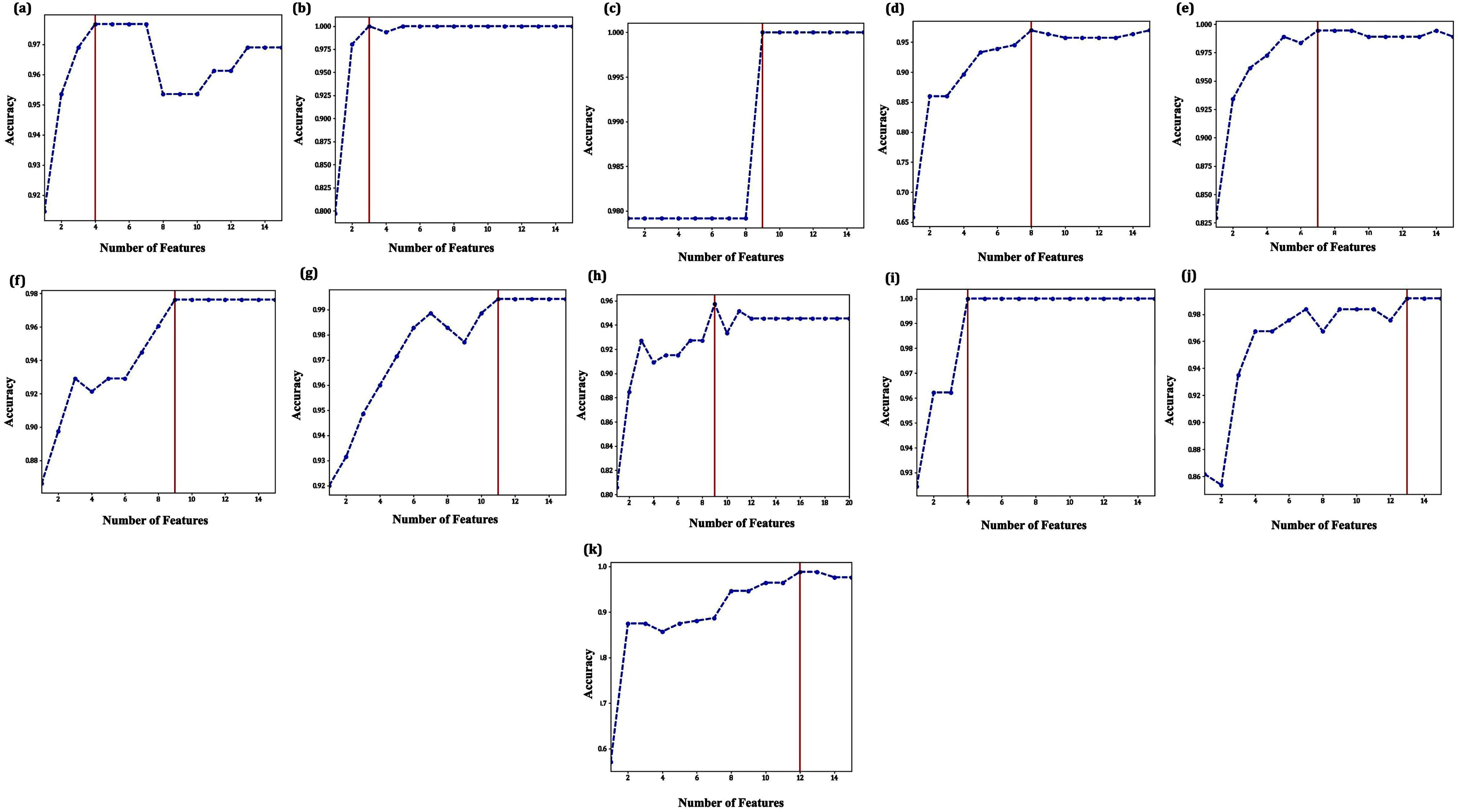
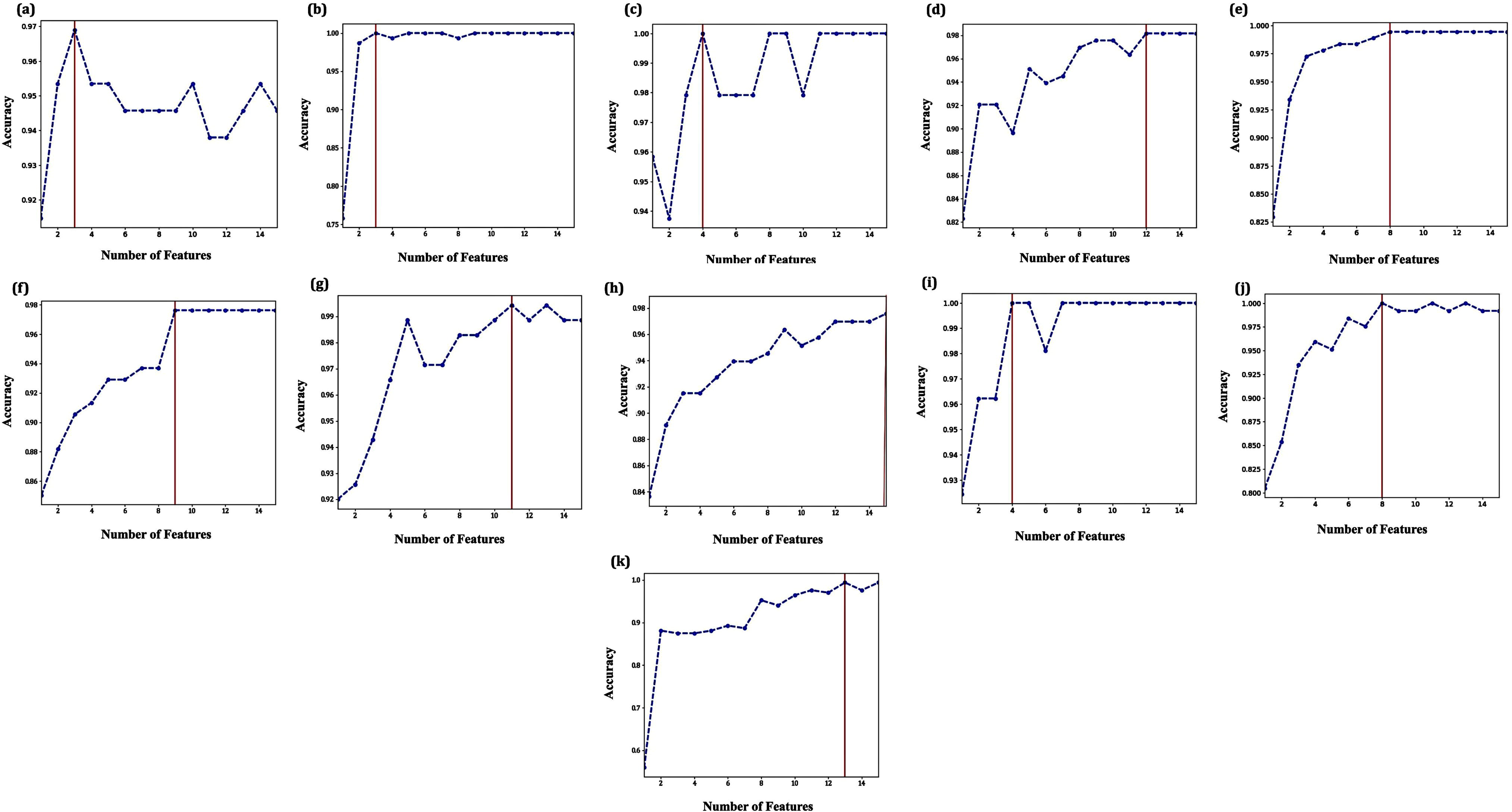
The test accuracy when the first 15 features with the higher fisher scores were applied as input features in (A) CSVM and (B) QSVM for (a) BLCA, (b) COAD, (c) GBM, (d) HNSC, (e) KIRC, (f) LIHC, (g) LUAD, (h) PRAD, (i) READ, (j) STAD, and (k) THCA.

Table 3 shows the optimum selected features by CSVM and skqulacs-QSVM. They are similar for COAD, LIHC, LUAD, READ, and are shared to some extent for other cancers.

**Table 3.**
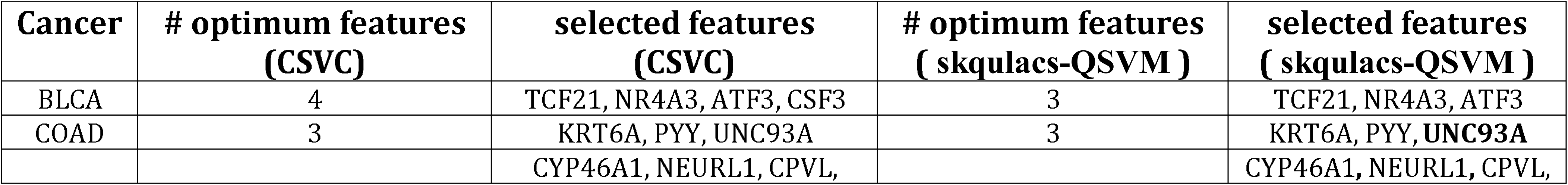
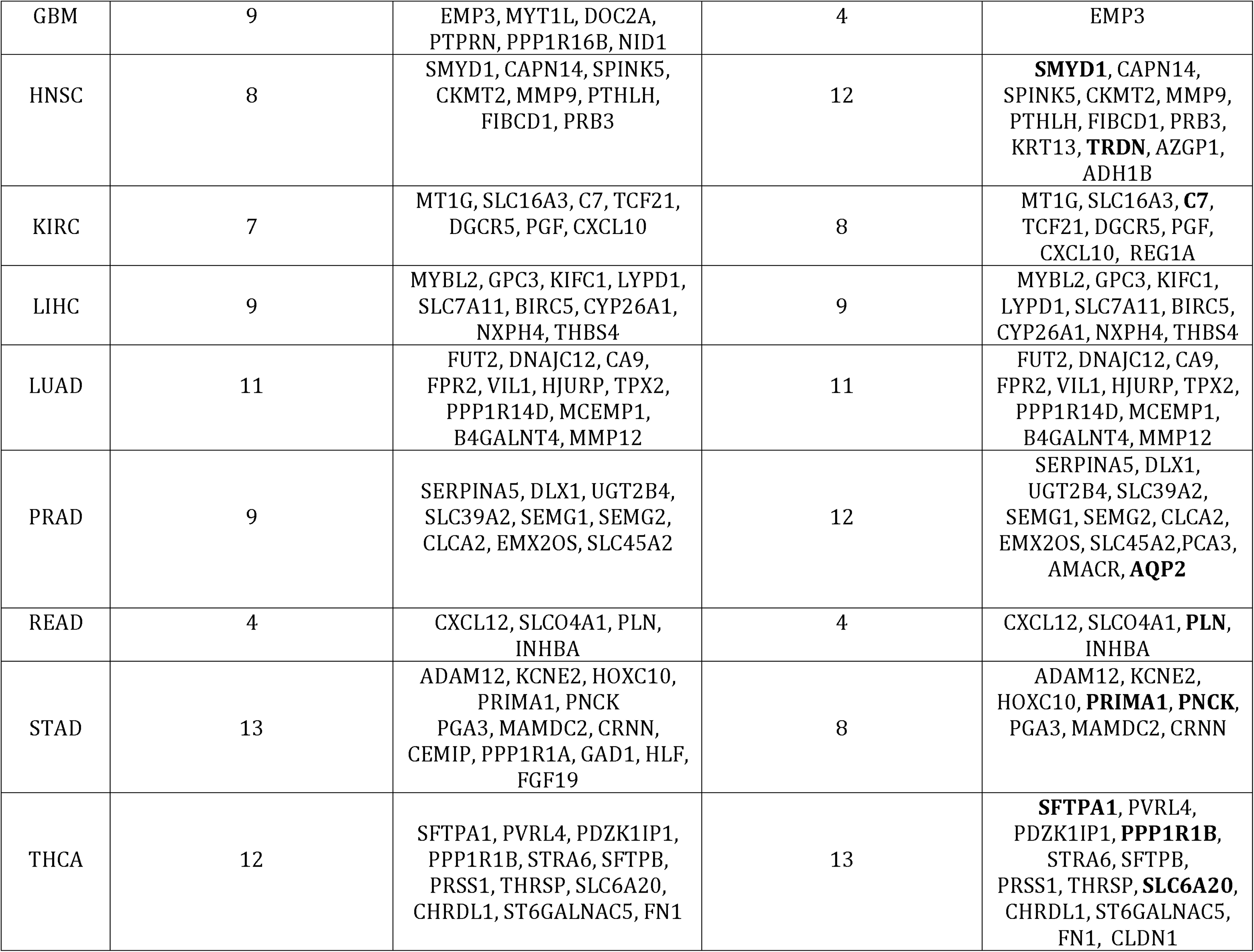
Number and the name of optimum features proposed as biomarkers.

Moreover, Figure 3 demonstrates the constructed circuits for all 11 cancers. We used parameterized quantum circuit (POC) structure to demonstrate the quantum circuit for each cancer data. In PQC, we used YZ-CX circuit constructions consisting of single-qubit y and z rotations and two-qubit entangling gates. Each quantum circuit is defined with N qubits that are used for implementing quantum machine learning algorithms. Depending on the cost of applying each element, we defined every circuit with layers that show the repetition of the circuit’s basic block. Here, for engineering the performance of quantum circuits for cancers with the lowest depth, we consider only one layer, *l* = 1, for each circuit. Our selection for circuits with one layer leads to have the best accuracy with respect to the construction elements (41).

**Figure 3:**
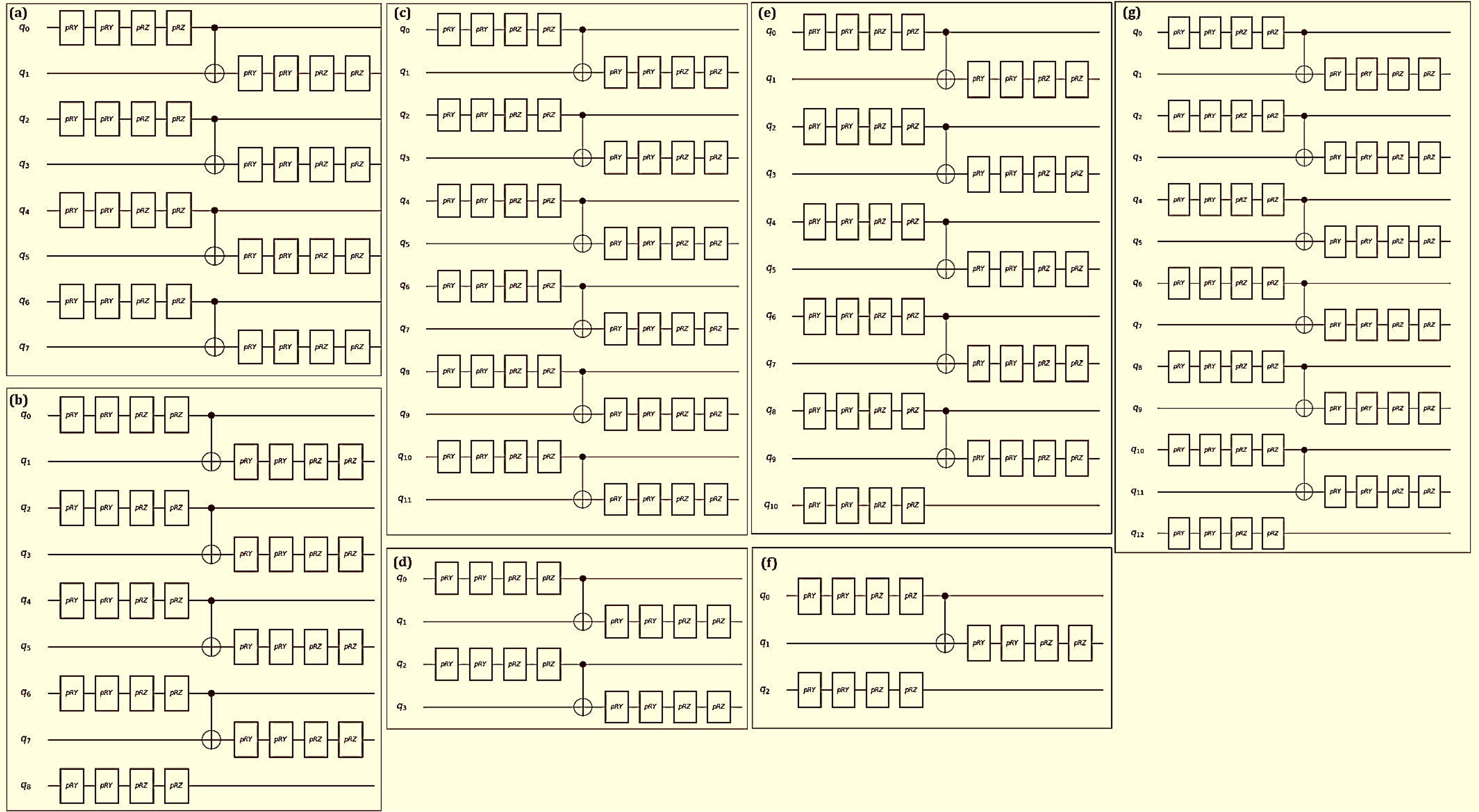
Schematic diagram of the parameterized quantum circuit for N qubits defined with only one layer. YZ-CX circuits consists of single-qubit y and z rotations and two-qubit entangling gates (a) N=8 for KIRC and STAD, (b) N=9 for LIHC, (c) N=12 for HNSC and PRAD, (d) N=4 for GBM and READ, (e) N=11 for LUAD, (f) N=3 for BLCA and COAD, (g) N=13 for THCA.

The classification reports including precision, recall, and F1-score as well as confusion matrices after constructing ML models show the advantages of skqulacs-QSVM over CSVM in most cancers (Figure 4). Across almost all eleven cancers, skqulacs-QSVM has yielded the best or similar classification outcomes.

**Figure 4:**
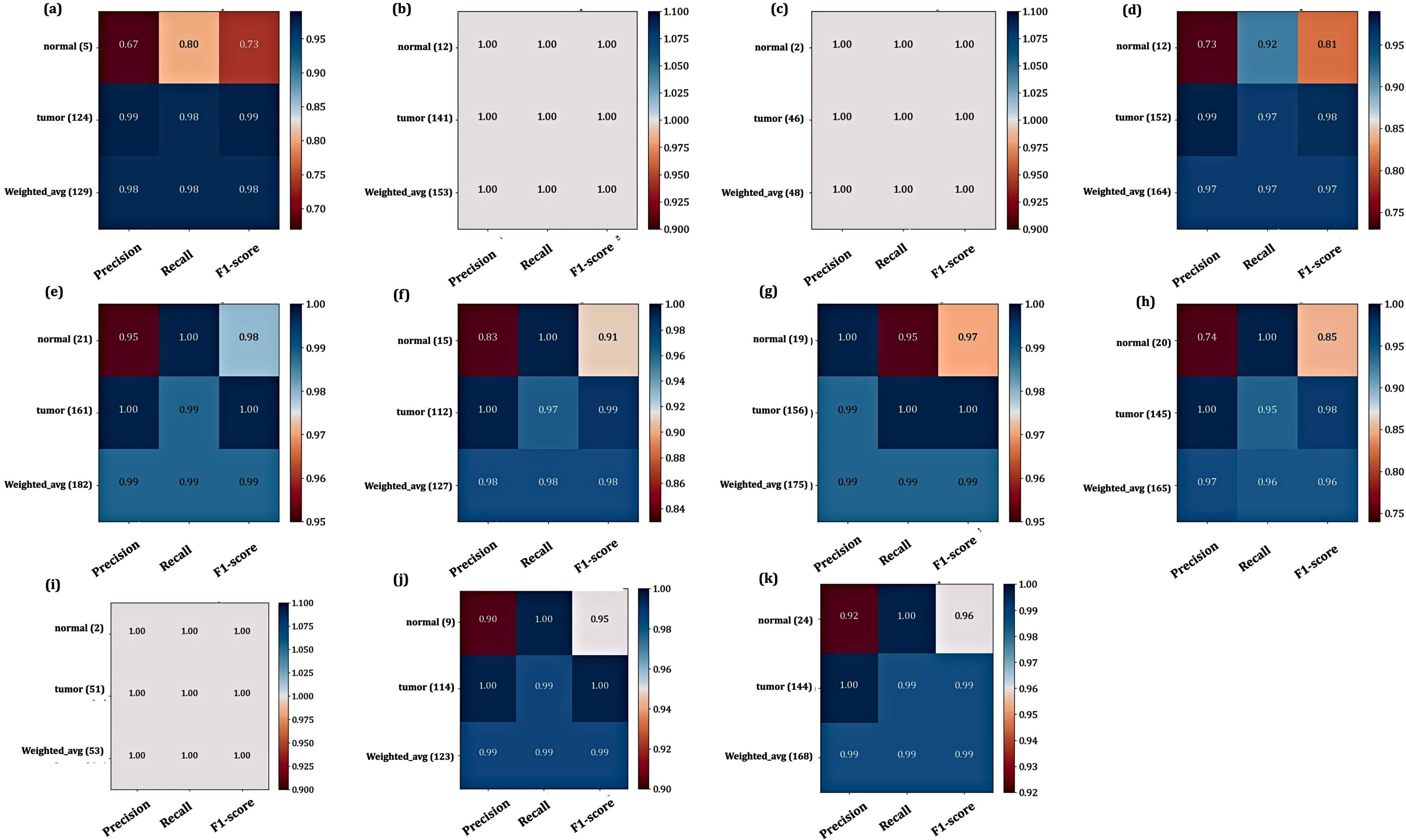
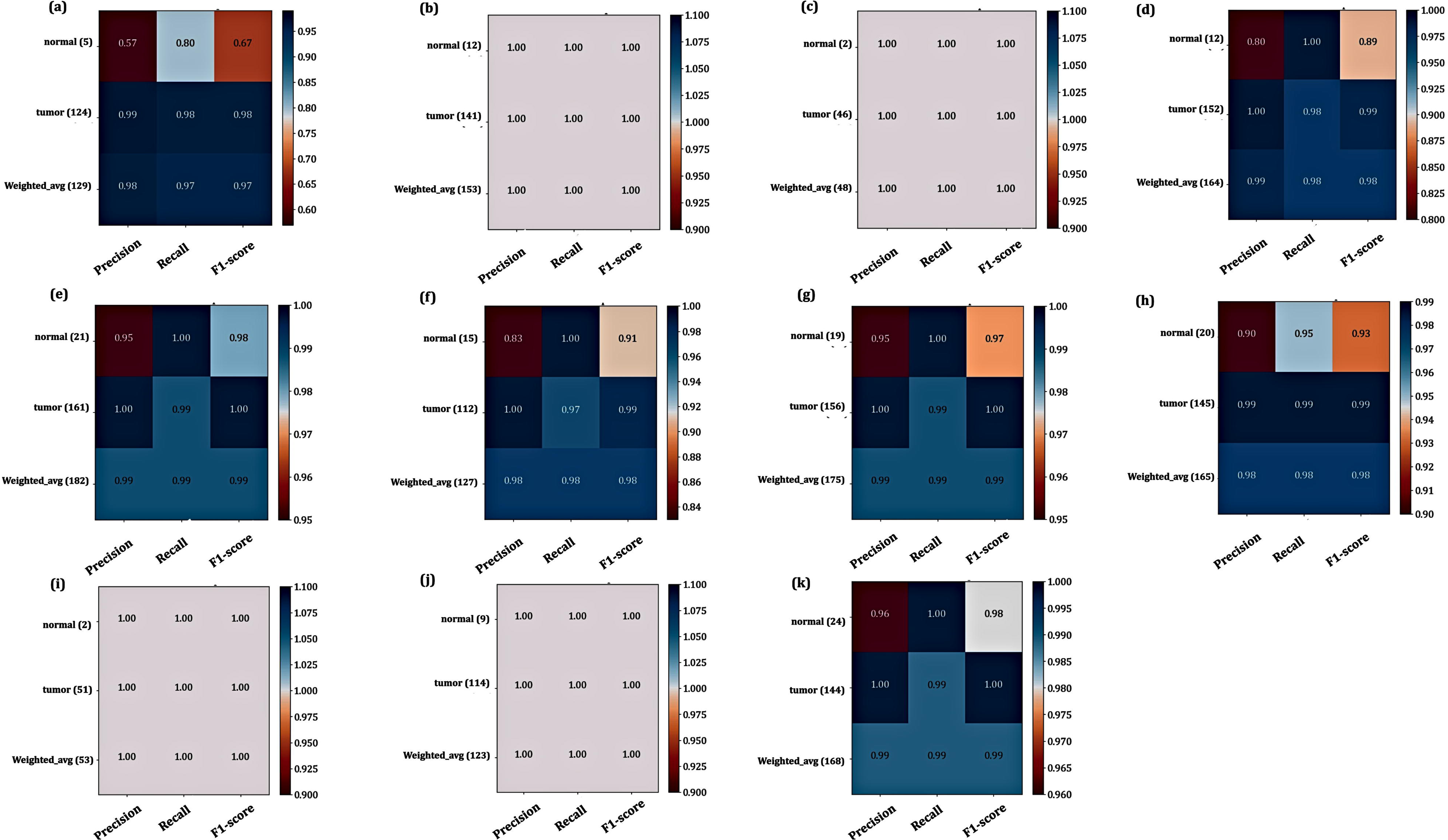
The classification reports including precision, recall, and F1-score for (a) BLCA, (b) COAD, (c) GBM, (d) HNSC, (e) KIRC, (f) LIHC, (g) LUAD, (h) PRAD, (i) READ, (j) STAD, and (k) THCA obtained by (A) CSVM and (B) QSVM.

### 4.3 Survival analysis

The heatmap of the expression of classifier genes for each cancer was drawn in all datasets of TCGA employing the UALCAN website (https://ualcan.path.uab.edu/analysis.html) (Figure 5) (42). It shows the difference in the expression of genes among normal and tumor samples. In addition, Kaplan-Meier plots were generated to show the effect of classifier gene expression on patient survival. The results revealed the remarkable effect of the following genes on the survival of patients: lower expressions of AZGP1, CKMT2, and SPINK5 in HNSC, higher expressions of BIRC5, KIFC1, MYBL2, and SLC7A11 in LIHC, lower expressions of MT1G and TCF21 as well as higher expression of PGF in KIRC, lower expression of MCEMP1 and higher expressions of HJURP and TPX2 in LUAD, lower expression of PLN in READ, higher expression of ADAM12 as well as lower expressions of MAMDC2 and PNCK in STAD (Figure S3-S8). Other genes had no effect on the patient’s survival.

**Figure 5:**
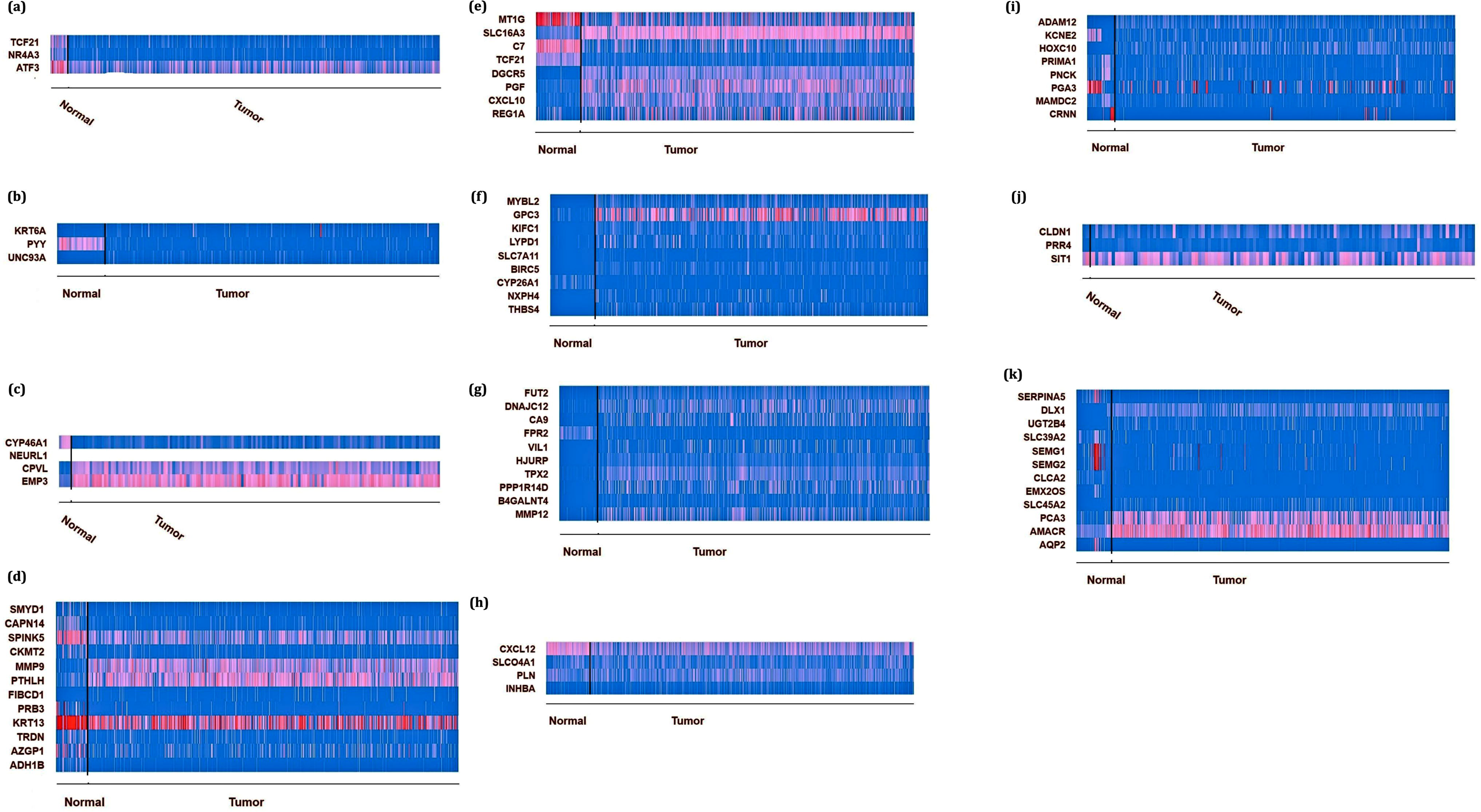
The heatmap of the expression of classifier genes for (a) BLCA, (b) COAD, (c) GBM, (d) HNSC, (e) KIRC, (f) LIHC, (g) LUAD, (h) PRAD, (i) READ, (j) STAD, and (k) THCA.

### 4.4 Promoter DNA methylation status and feature’s expression

DNA promoter methylation is an epigenetic modification that implicates the regulation of gene expression and often happens in the early phase of tumorigenesis (43). We explored the promoter DNA methylation status of the classifier genes by producing the box-whisker plots (42). We used Welch’s T-test that was to assess the meaningfulness of differences in expression levels between normal and tumor groups. Figure S9-S17 demonstrates the methylation level of features that had an opposite relationship with the expression value (44). As shown, the following changes in the promotor DNA methylation might be the conceivable mechanisms leading to the downregulation or upregulation of classifier gene expression obtained by skqulacs-QSVM: significant declined promotor DNA methylation levels of KRT6A and UNC93A in COAD, MMP9 and PTHLH in HNSC, REG1A and SLC16A3 in KIRC, MYBL2, GPC3, KIFC1, LYPD1, SLC7A11, and BIRC5 in LIHC, FUT2, CA9, VIL1, TPX2, PPP1R14D, B4GALNT4, and MMP12 in LUAD, UGT2B4, AMACR, and SLC45A2 in PRAD, INHBA and CXCL12 in READ, SFTPA1, PVRL4, PPP1R1B, STRA6, SFTPB, PDZK1IP1, THRSP, FN1, and CLDN1 in THCA accompany with increased gene expression as well as significantly increased promotor DNA methylation level of TCF21 in BLCA, CKMT2 and KRT13 in HNSC, TCF21 and CXCL10 in KIRC, CYP26A1 in LIHC, FPR2 in LUAD, EMX2OS and AQP2 in PRAD.

### 4.5 Pathway enrichment analysis

The pathway enrichment revealed that some of the identified genes by CSVM and skqulacs- QSVM are significantly involved in known pathways activated in various cancers (Figure 6).

**Figure 6:**
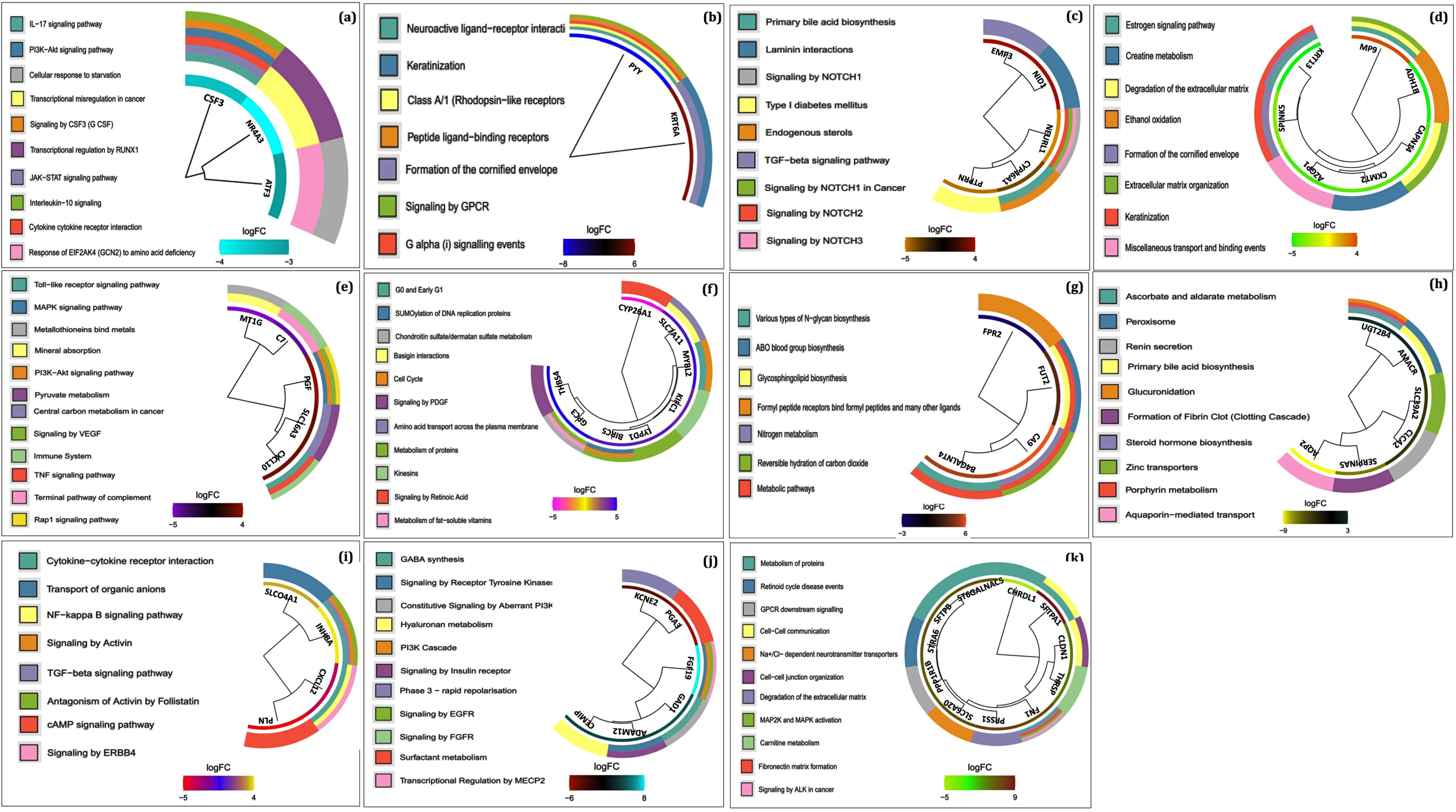
The enriched pathways by the classifier genes of (a) BLCA, (b) COAD, (c) GBM, (d) HNSC, (e) KIRC, (f) LIHC, (g) LUAD, (h) PRAD, (i) READ, (j) STAD, and (k) THCA.

## 5. Discussion

A quantum kernel is a function determining the resemblance between two quantum states in the feature space (24). Employing kernel, the QML algorithms classify quantum states according to their similarities. The infinite possibility of the dimension of the kernel Hilbert space makes the kernel approach powerful (23).

In this paper, we surveyed the potential of QML for the classification of eleven cancers based on the RNA-seq gene expression data. To this end, we applied the hybrid classical-quantum machine learning algorithm and constructed quantum circuits and then skqulacs-QSVM models for each cancer. The outcomes revealed the remarkable classification metrics for skqulacs- QSVM.

There are few reports regarding the application of quantum models for the classification of cancers. A quantum annealer, known as D-Wave 2000Q, was employed to construct a model based on gene expression data to predict if non-small cell lung cancer patients have either squamous cell carcinoma or adenocarcinoma (45). The univariate statistics and XGBoost were used as the feature selection approaches. In this research, the performance of quantum machine learning was reported as significant and robust. In another study, a hybrid quantum-mechanical system with 2 qubits operation was used to encode, process, and classify images of cancerous and non-cancerous pigmented skin lesions (46). This approach dominated the constraints of classical computing models. The QSVM model was also utilized to unravel the classification of malignant breast cancer tumors from non-cancerous benign tumors (47). At first, the principal component analysis algorithm was applied to decrease the feature’s number based on the accessible number of qubits. The proposed model could resolve the binary task of breast cancer diagnosis in logarithmic order contrary to the conventional polynomic approach leading to speedy classification. The main restriction of this study is using only two principal components (2 features) that could just explain up to 70% of the variance in data.

For the first time, we proposed a stepwise method to show the power of QML in classifying various cancers from healthy subjects. In addition, we could determine several features as biomarkers. These selected biomarkers have significant functions in developing cancer by involvement in regulating several cancer-related pathways. Some of these classifier genes affect the survival of cancer patients. In addition, some of them have dysregulated promoter DNA methylation that might impress the expression of genes. We performed text mining to find the previously reported gene/protein biomarkers among the classifiers. Supplementary file 3 contains the published papers that reported the roles of the identified features in each cancer. It has been revealed that many of them have been introduced as diagnostic/prognostic biomarkers. Such results confirm the capability of skqulacs-QSVM in determining cancer biomarkers. On the other hand, some novel biomarkers were also determined which are indicated as bold in Table 3. In the following, we discuss the novel identified biomarkers.

UNC93A was found a novel gene involved in COAD. The highly polymorphic and truncating mutations in UNC93A have been observed in ovarian tumors and cell lines. Polymorphic variants of genes in individuals may increase the risk of cancer (48). It was overexpressed and hypomethylated in COAD samples in our study. Therefore, it may promote the progression of this cancer.

SMYD1 and TRDN are two potential biomarkers for HNSC. SMYD1 is a specific protein for skeletal and cardiac muscles and targets histone 3, lysine 4 (H3K4) methylation. It contributes to cell differentiation, regulation of embryonic development, and cardiomyocyte specification (49). SMYD1 is also involved in immunity through the IFNγ signaling pathway and is a downstream effector of IFNγ (50). It was downregulated in HNSC and might be involved in providing proper conditions for the development of HNSC. TRDN has a key role in normal skeletal muscle strength through its function in skeletal muscle excitation-contraction coupling as a segment of the calcium release complex. The interaction of TRDN interaction with CASQ2 and RYR2 regulates the release of RYR2-mediated calcium (51). The Underexpression of TRDN may disrupt calcium release. The unusual Ca2+ signaling might contribute to tumorigenesis in HNSC.

The C7 gene is the terminal part of the complement cascade, which was downregulated in KIRC. It could function as a tumor suppressor (52); therefore, the changes in its expression could accompany tumor progression.

Aquaporins (AQPs) are transmembrane water channel proteins that are mostly overexpressed in different human cancers and are involved in the development of prostate cancer after therapy. AQP4 and AQP7 have been determined as biomarkers for prostate cancer (53). From the results of our study, we also identified AQP2 as a feasible biomarker for PRAD.

PLN is negatively-regulates the Ca2+ uptake action of the energy-expending SR Ca2+ ATPase (SERCA2a). The unphosphorylated PLN attaches to SERCA2a and interdicts its function. After phosphorylation, the inhibitory interaction between PLN and SERCA2a is ended and the apparent Ca2+ affinity is increased. The disturbance in the SERCA activity by the downregulation of PLN might interrupt the Ca2+ homeostasis and promote apoptosis (54) in READ.

PRIMA1 and PNCK were determined to be involved in STAD. PRIMA1 encodes a membrane protein that binds acetylcholinesterase to cell membranes. Downregulation of PRIMA1 results in declined AChE activity, which in turn more cholinergic transmission would occur (55). The cholinergic anti-inflammatory pathway has a role in different inflammatory diseases. It has been hypothesized that the affirmative modulation of cholinergic neurotransmission may have a remarkable effect on stomach adenocarcinoma through the parasympathetic signaling/cholinergic anti-inflammatory pathway (56). PNCK is Calcium/Calmodulin-Dependent Protein Kinase Type 1B and belongs to a member of the CaM kinase I family. It possesses a substantial duty in human malignancies and likely in STAD. PNCK could have different functions in various cancers (57). The downregulation of PNCK could inhibit tumor proliferation and induce apoptosis in nasopharyngeal carcinoma (58). However, the underlying mechanism of the involvement of PNCK has not been fully determined.

In THCA, three genes including SFTPA1, PPP1R1B, and SLC6A20 have been identified as possible diagnostic biomarkers. SFTPA1 belongs to the C-type lectin family, which plays a major role in retaining the homeostasis of lung tissue and regulation of the innate immune function via the TLR signaling pathway in lung cancer (59). Therefore, its upregulation in THCA may involve in promoting cancer. PPP1R1B encodes a bifunctional signal transduction molecule. The function of this molecule as a phosphatase or kinase inhibitor is regulated by dopaminergic and glutamatergic receptor stimulation. The overexpression of PPP1R1B in breast, colon, and gastric cancer, as well as THCA in this study, have been detected. PPP1R1B may control pro-oncogenic signal transduction pathways to elevate cancer cell viability (60, 61). SLC6A20 is a protein-coding gene that acts as a proline transporter expressed in the kidney and small intestine. Similar to THCA in our study, the upregulation of SLC6A20 in various malignancies has been observed in many human pan-cancer samples (62).

## 6. Conclusion

In this work, we presented a hybrid classical-quantum machine learning algorithm for classifying RNA-seq cancer data. The outcomes confirmed the capability of quantum machine learning in the analysis of high-throughput real-world biological data. The consistency of the QML results with CML shows that we could profit from the advantage of QML such as exponential speed-up for the analysis of high-throughput real-world biological data.

## Availability of data and materials

The RNA-seq data expression were downloaded from UCSC Xena database (https://xenabrowser.net/). The codes are available in https://github.com/Mohadesehzarei/skqulacs_QSVM.

