## Supplementary Data Files for "Quantum machine learning for untangling the real-world problem of cancers classification based on gene expressions": Supplementary figures.docx

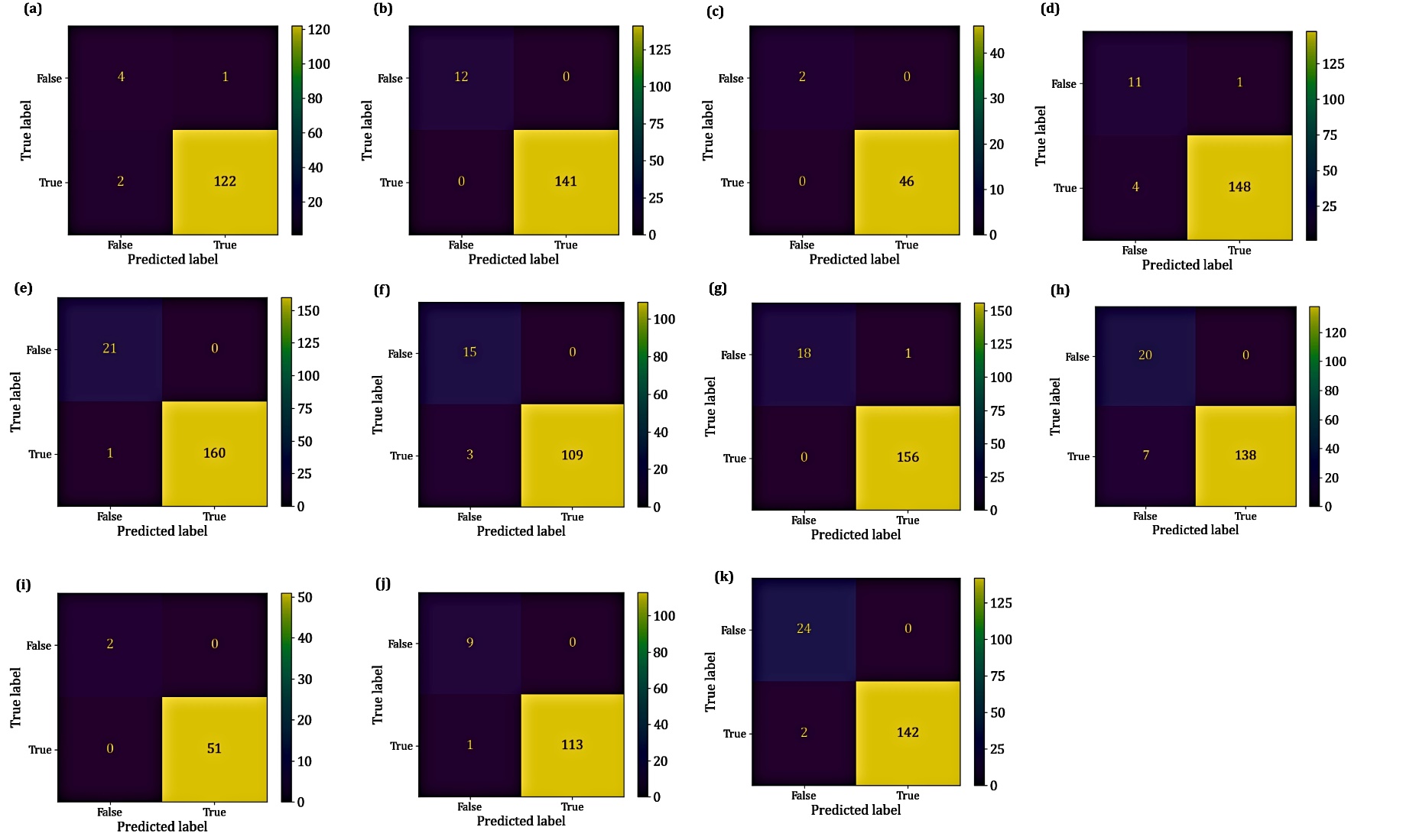

Figure S1. Confusion matrix for test set after implementation CSVM for a) BLCA, (b) COAD, (c) GBM, (d) HNSC, (e) KIRC, (f) LIHC, (g) LUAD, (h) PRAD, (i) READ, (j) STAD, and (k) THCA.

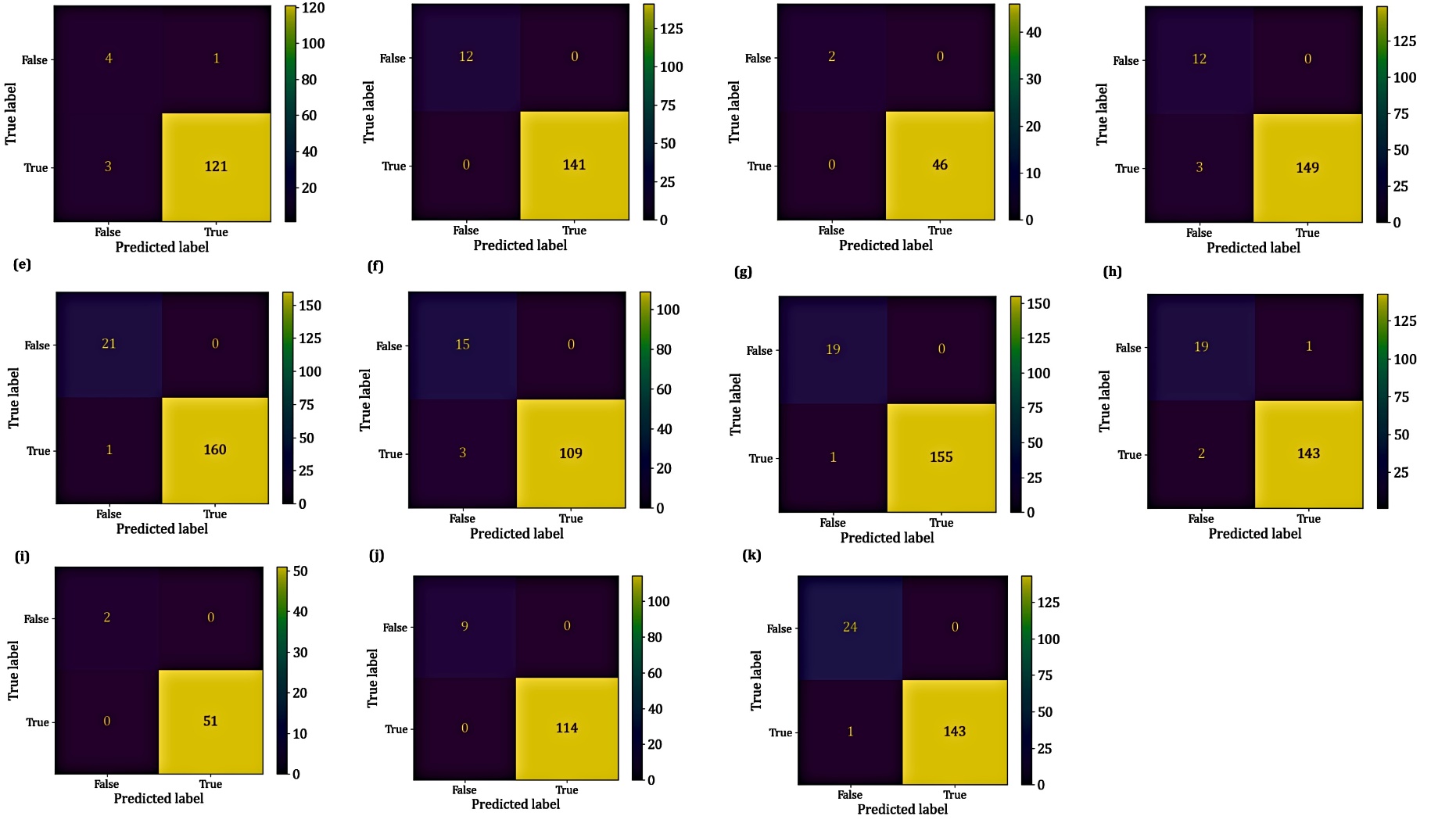

**Figure S2.** Confusion matrix for test set after implementation QSVM for a) BLCA, (b) COAD, (c) GBM, (d) HNSC, (e) KIRC, (f) LIHC, (g) LUAD, (h) PRAD, (i) READ, (j) STAD, and (k) THCA.

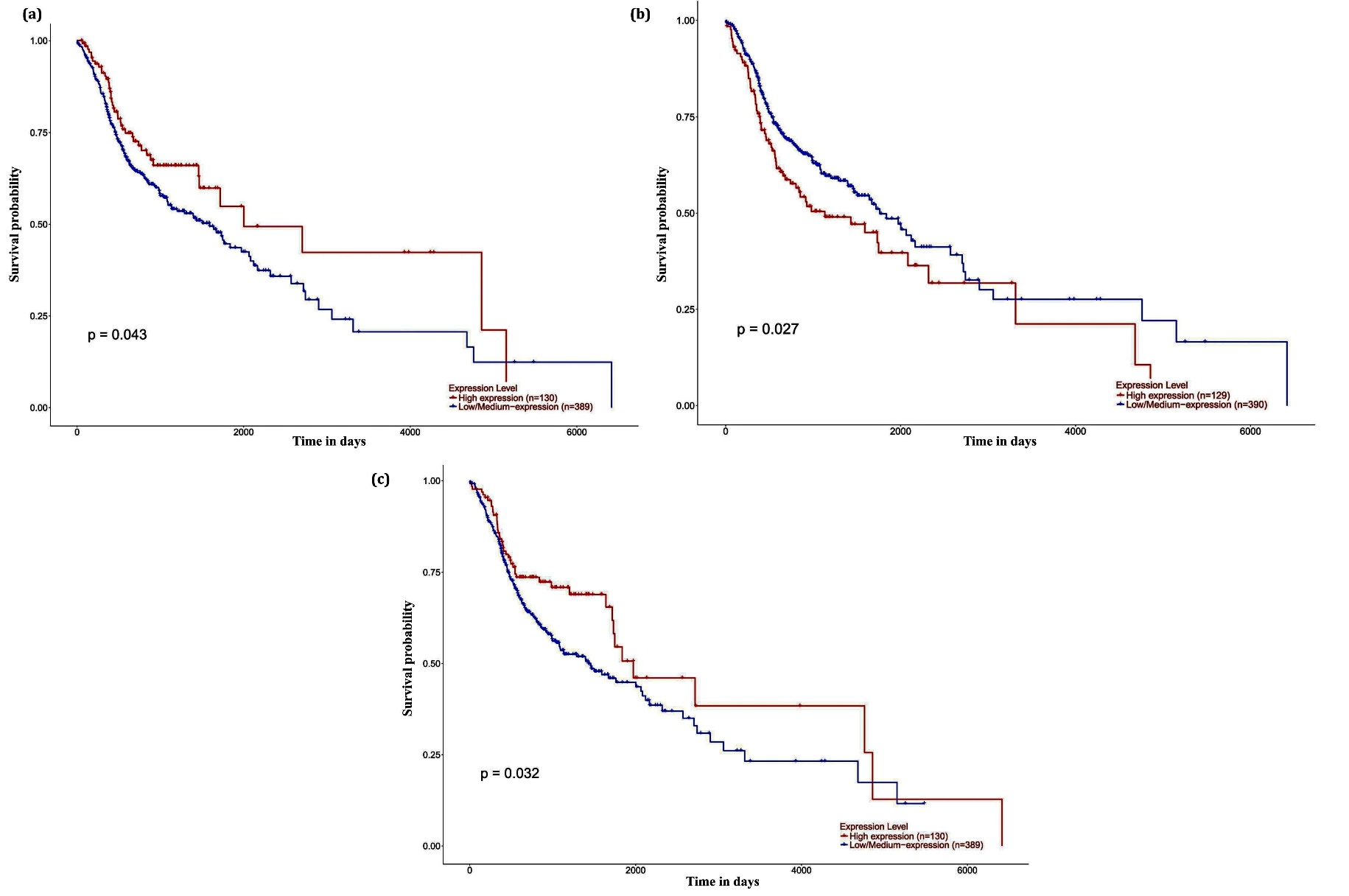

Figure S3. The effect of (a) AZGP1, (b) CKMT2, and (c) SPINK5 on the survival of HNSC patients.

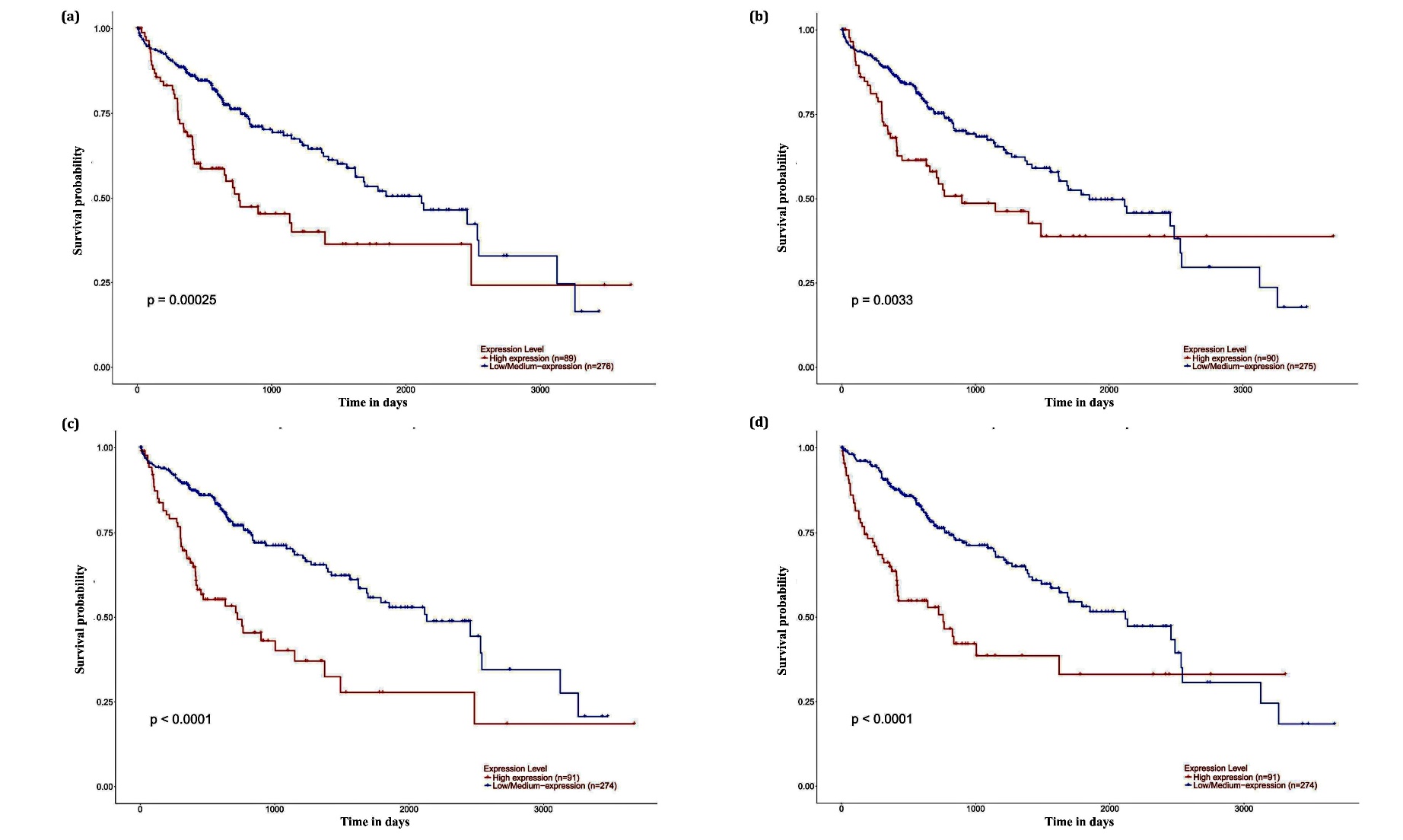

Figure S4. The effect of (a) BIRC5, (b) KIFC1, (c) MYBL2, and (d) SLC7A11 on the survival of LIHC patients.

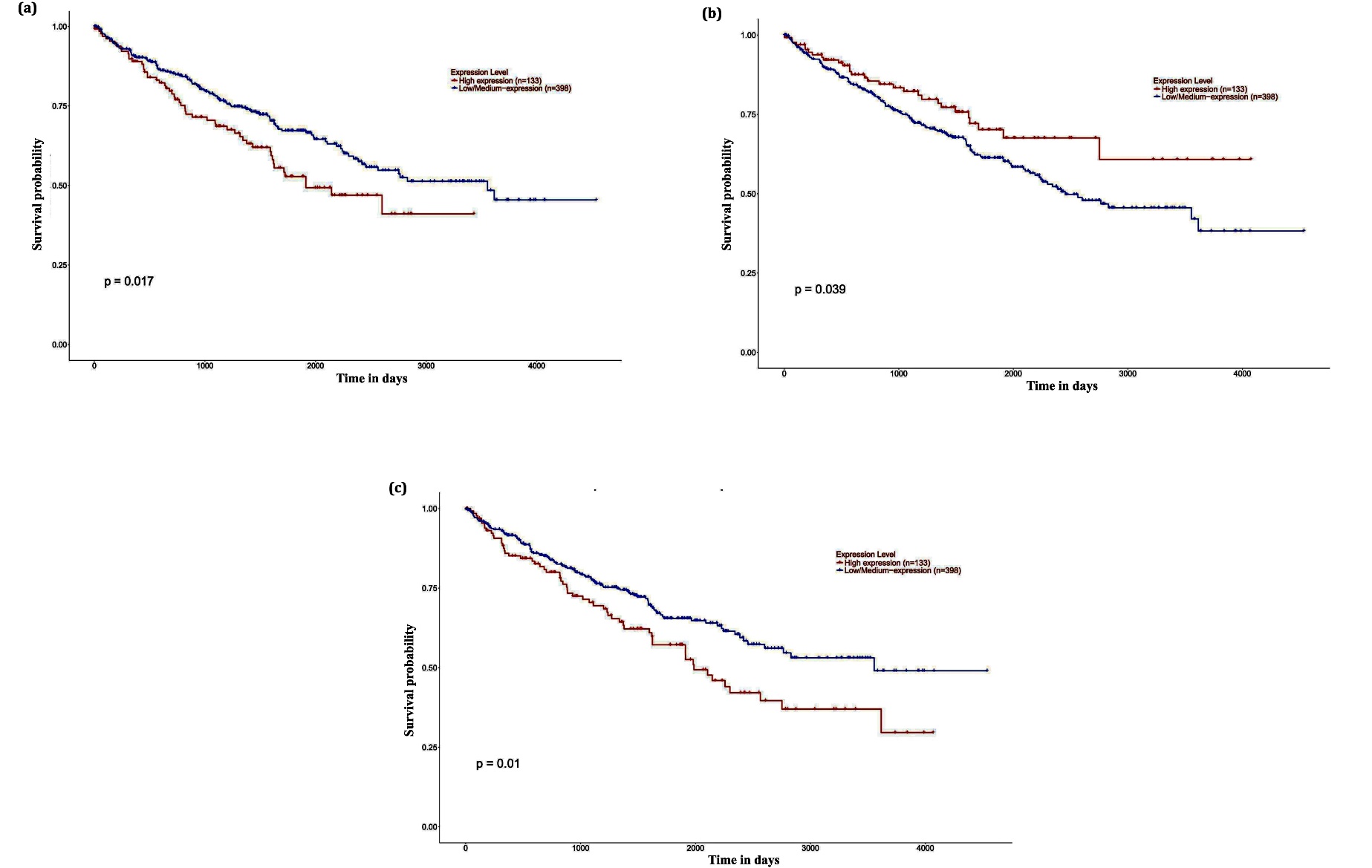

Figure S5. The effect of (a) MT1G, (b) TCF21, and (c) PGF on the survival of KIRC patients.

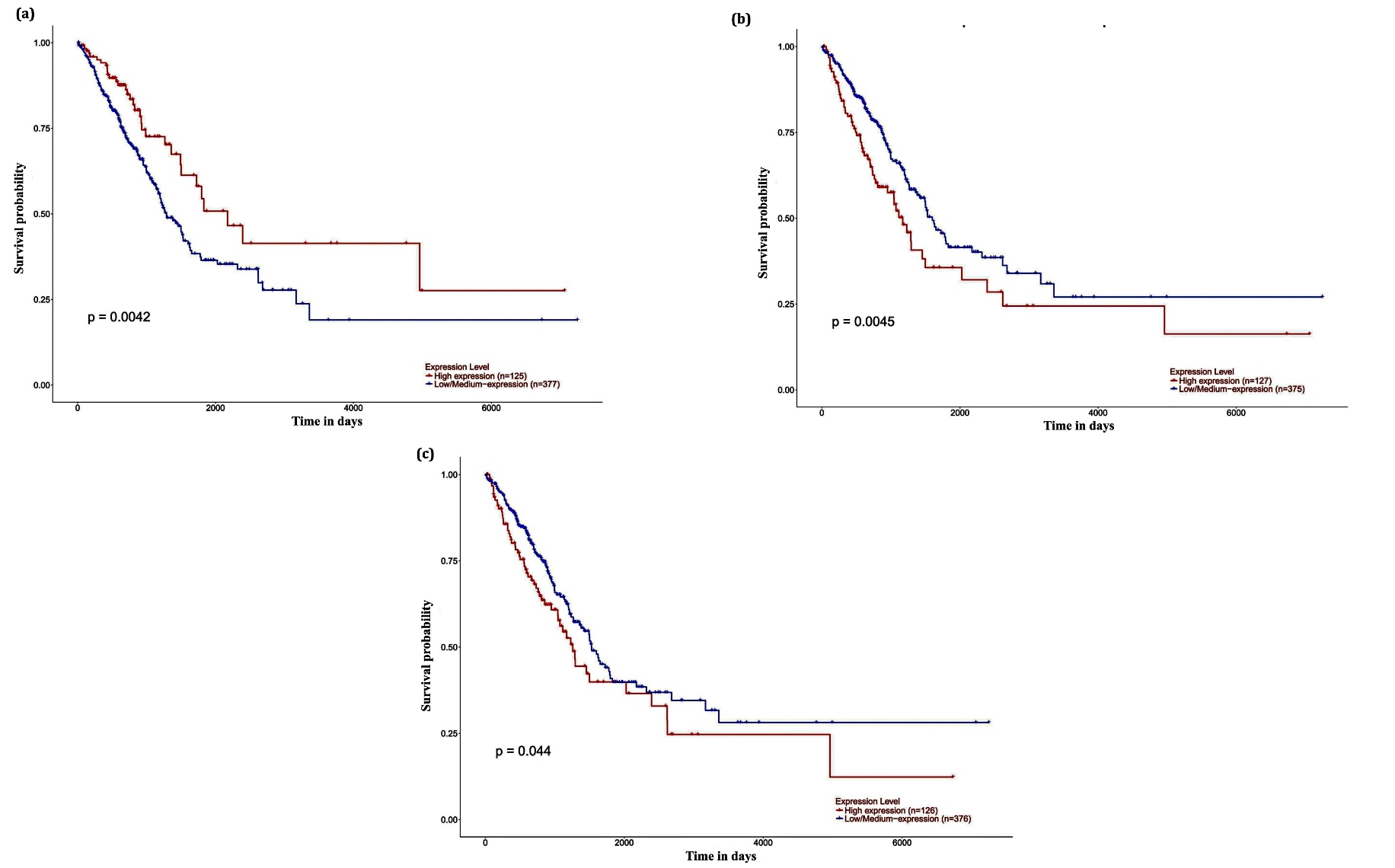

Figure S6. The effect of (a) MCEMP1, (b) HJURP, and (c) TPX2 on the survival of LUAD patients.

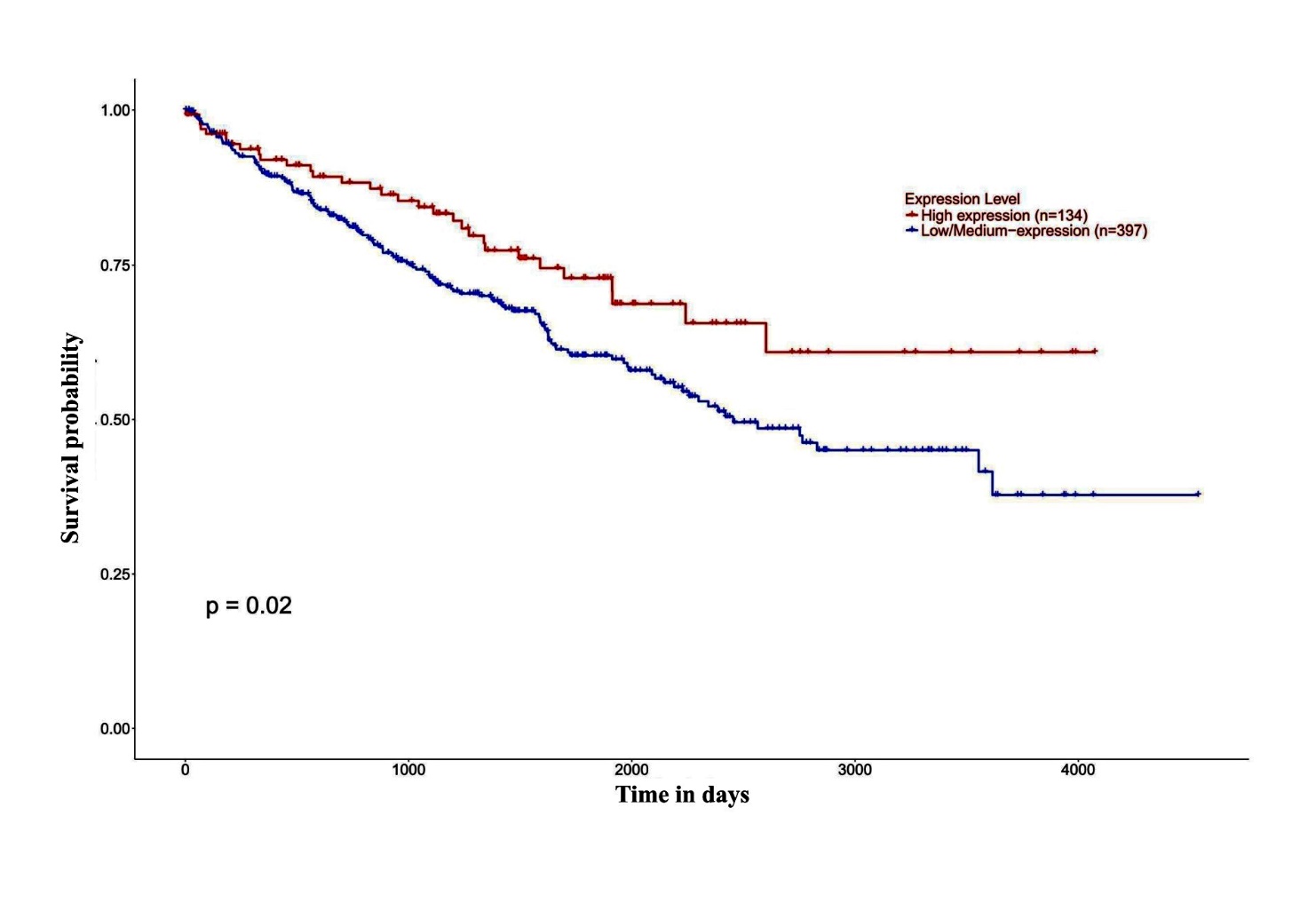

Figure S7. The effect of PLN on the survival of READ patients.

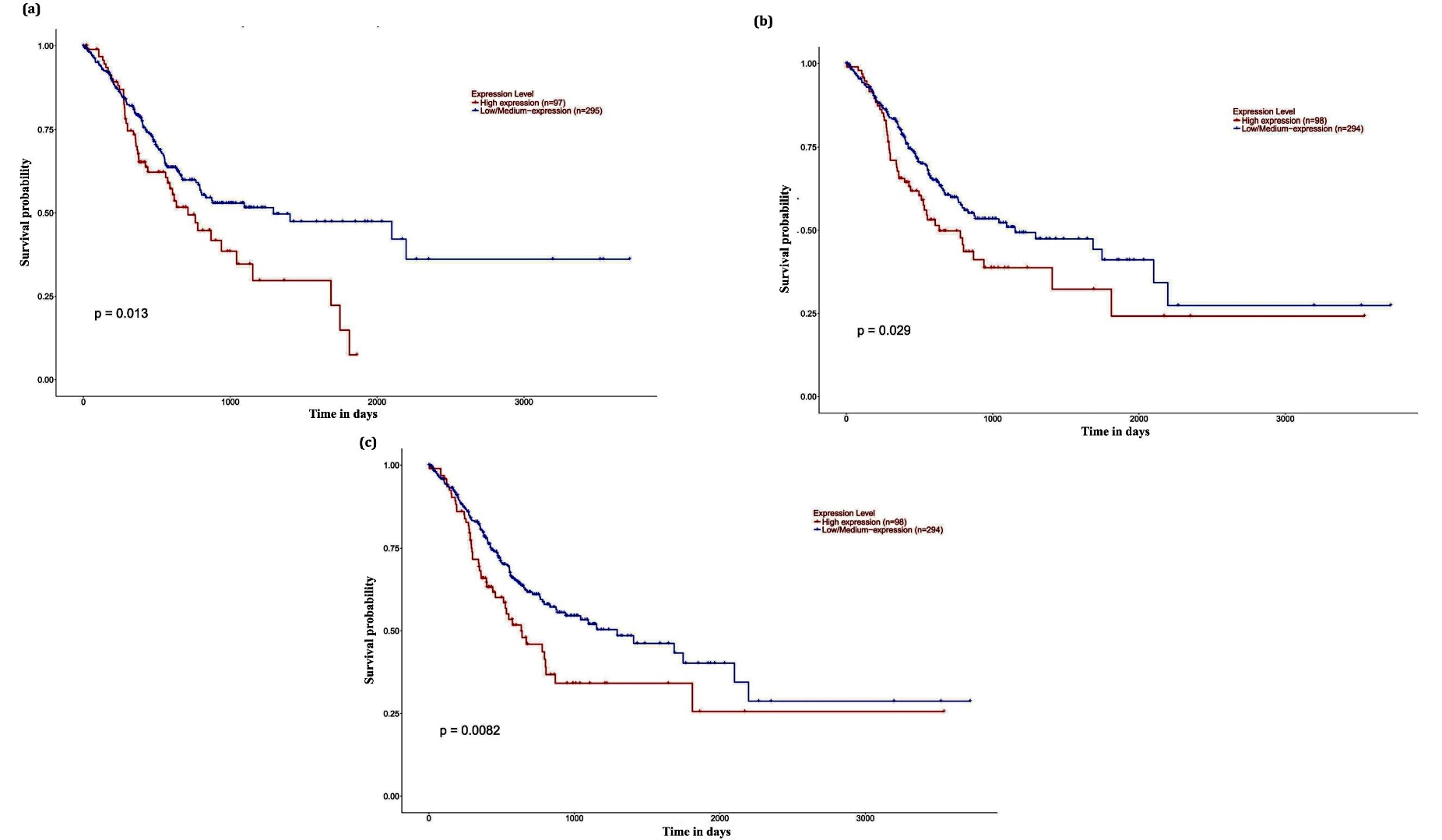

Figure S8. The effect of (a) ADAM12, (b) MAMDC2, and (c) PNCK on the survival of STAD patients.

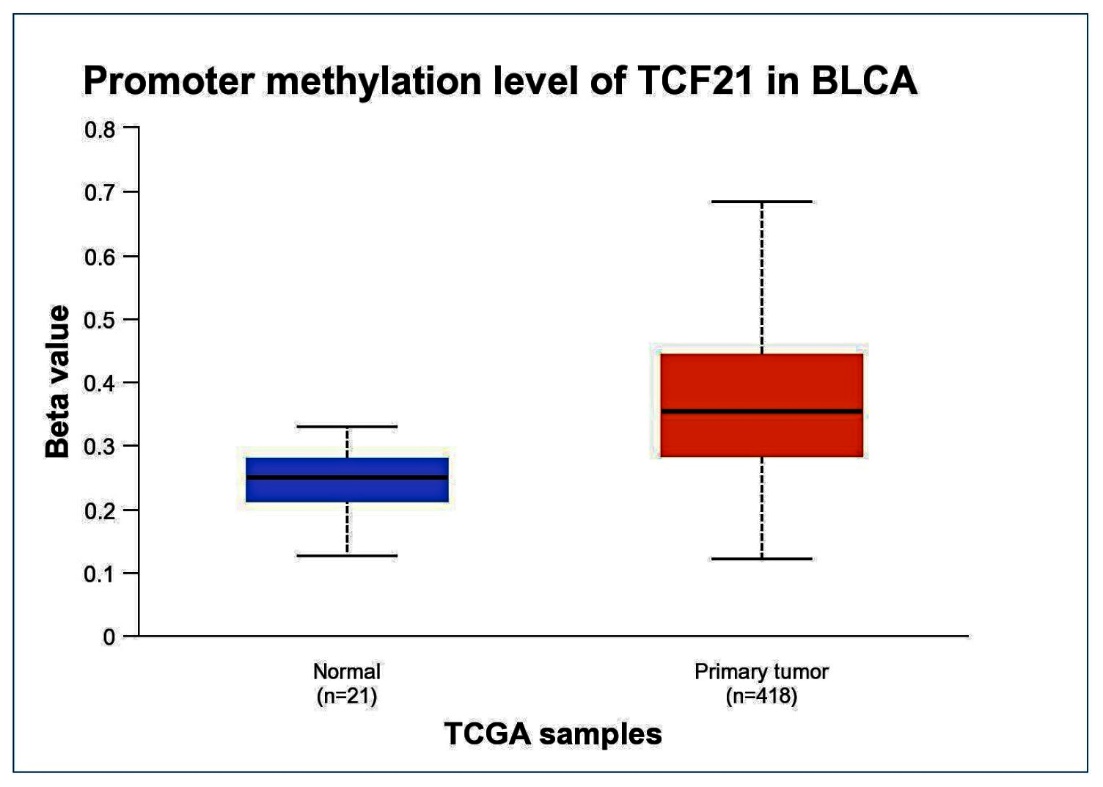

Figure S9. The significant change in promotor DNA methylation levels of TCF21 in BLCA.

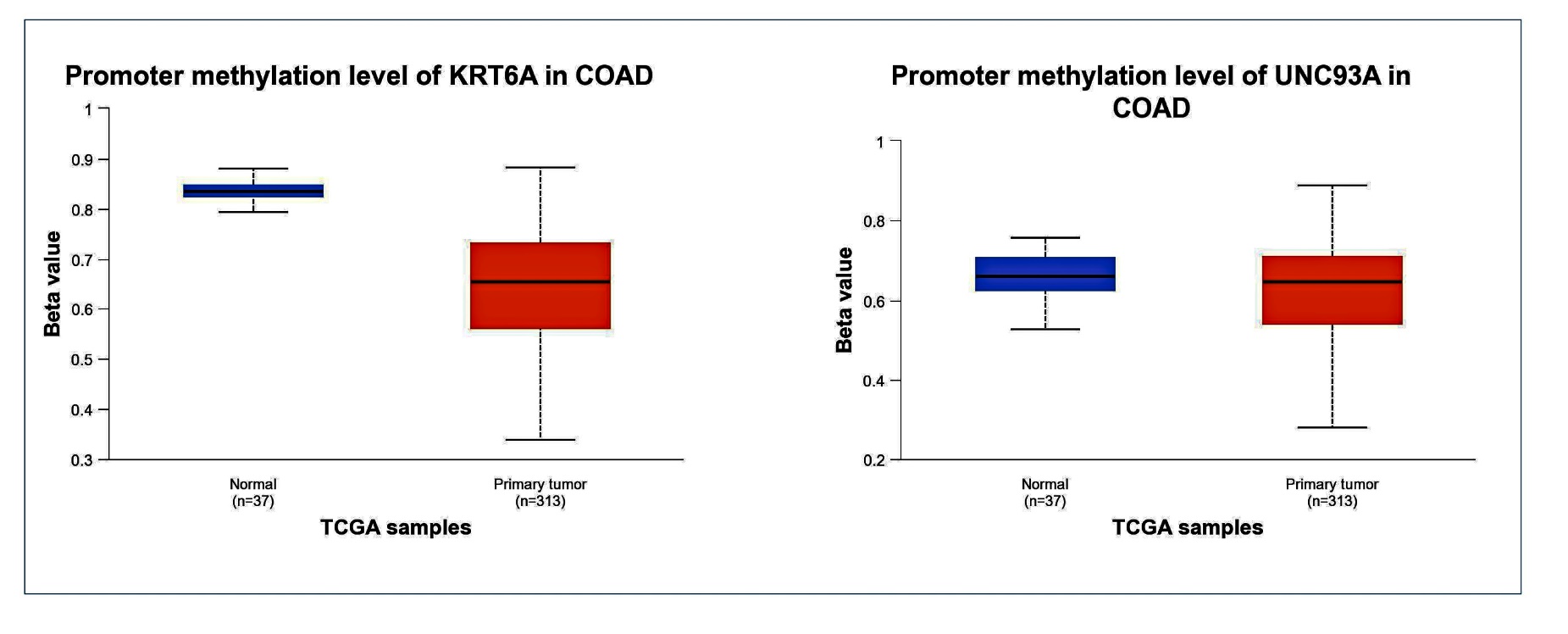

Figure S10. The significant changes in promotor DNA methylation levels of KRT6A and UNC93A in COAD.

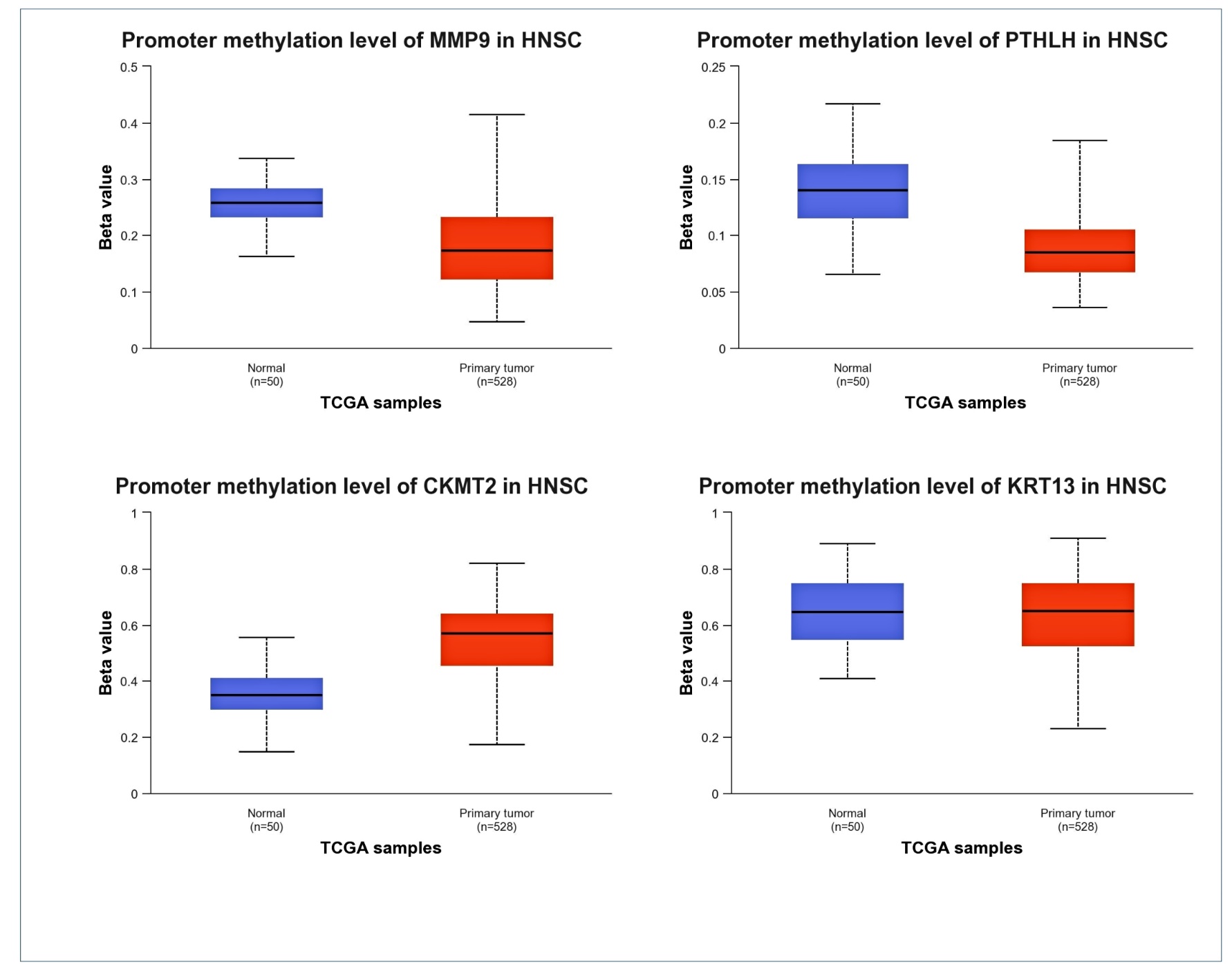

Figure S11. The significant changes in promotor DNA methylation levels MMP9, PTHLH, CKMT2, and KRT13 in HNSC.

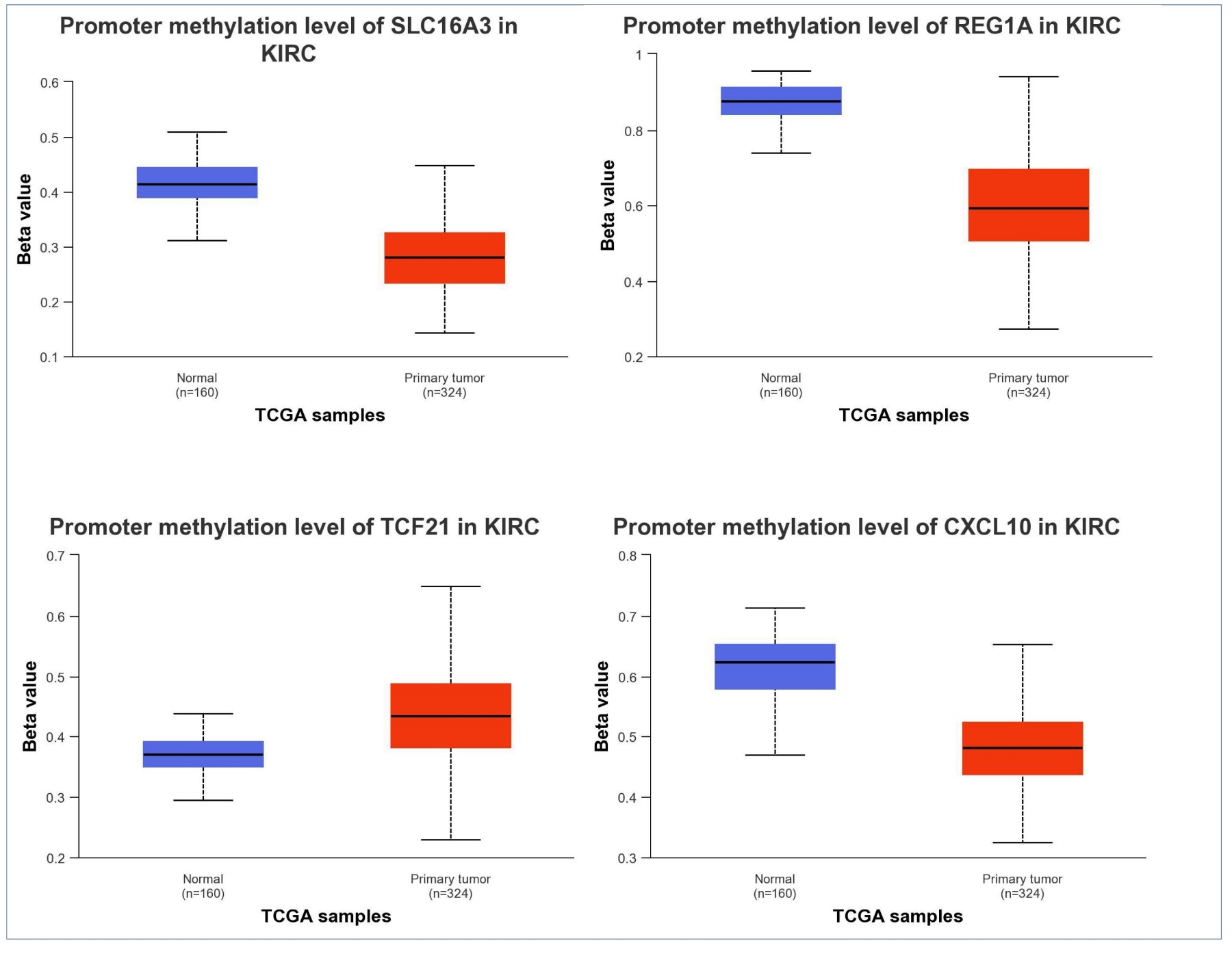

Figure S12. The significant change in promotor DNA methylation levels of SLC16A3, REG1A, CXCL10, and TCF21 in KIRC.

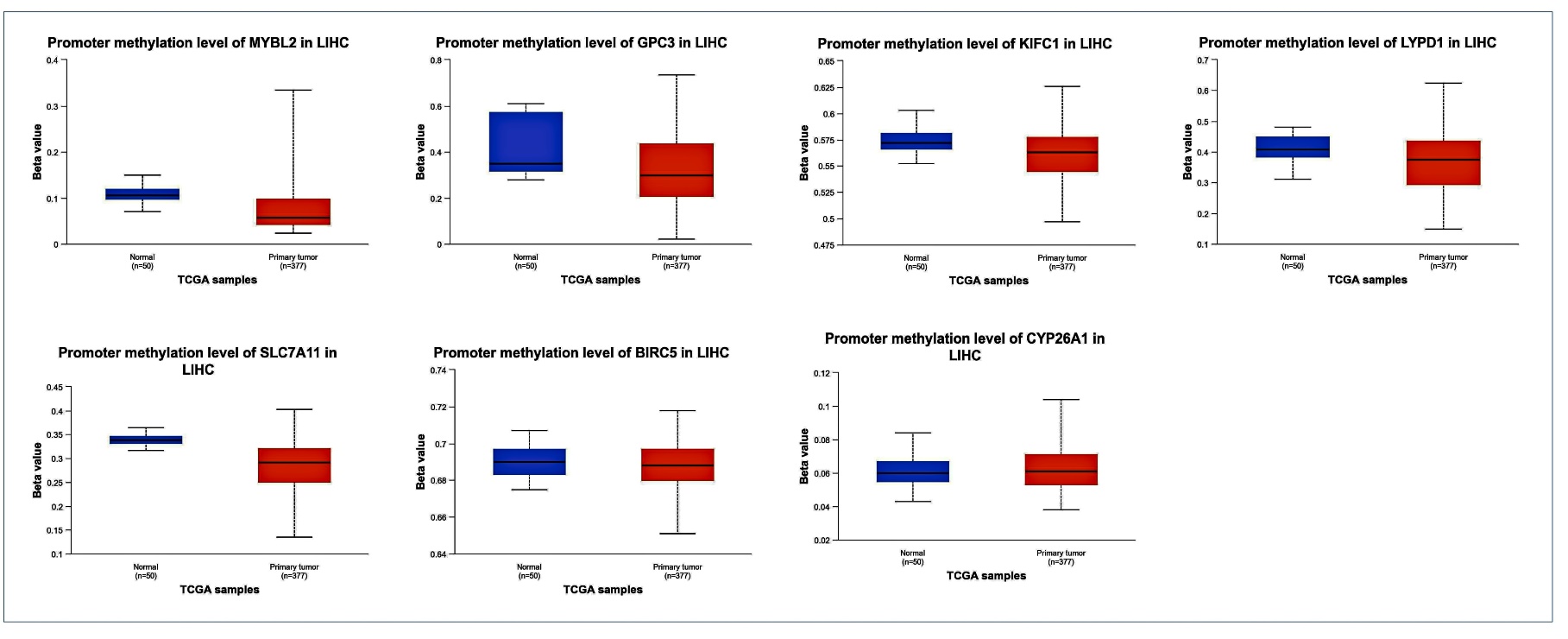

Figure S13. The significant change in promotor DNA methylation levels of MYBL2, GPC3, KIFC1, LYPD1, SLC7A11, CYP26A1, and BIRC5 in LIHC.

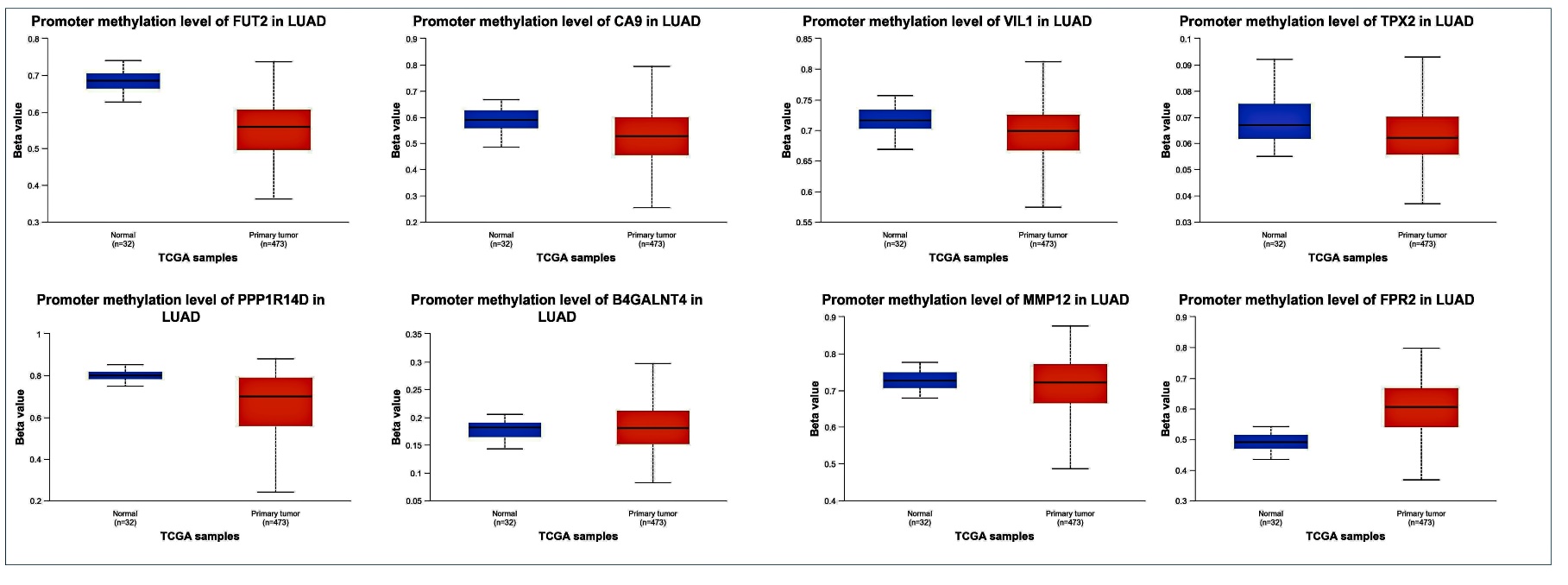

Figure S14. The significant change in promotor DNA methylation levels of FUT2, CA9, VIL1, TPX2, PPP1R14D, B4GALNT4, MMP12, and FPR2 in LUAD.

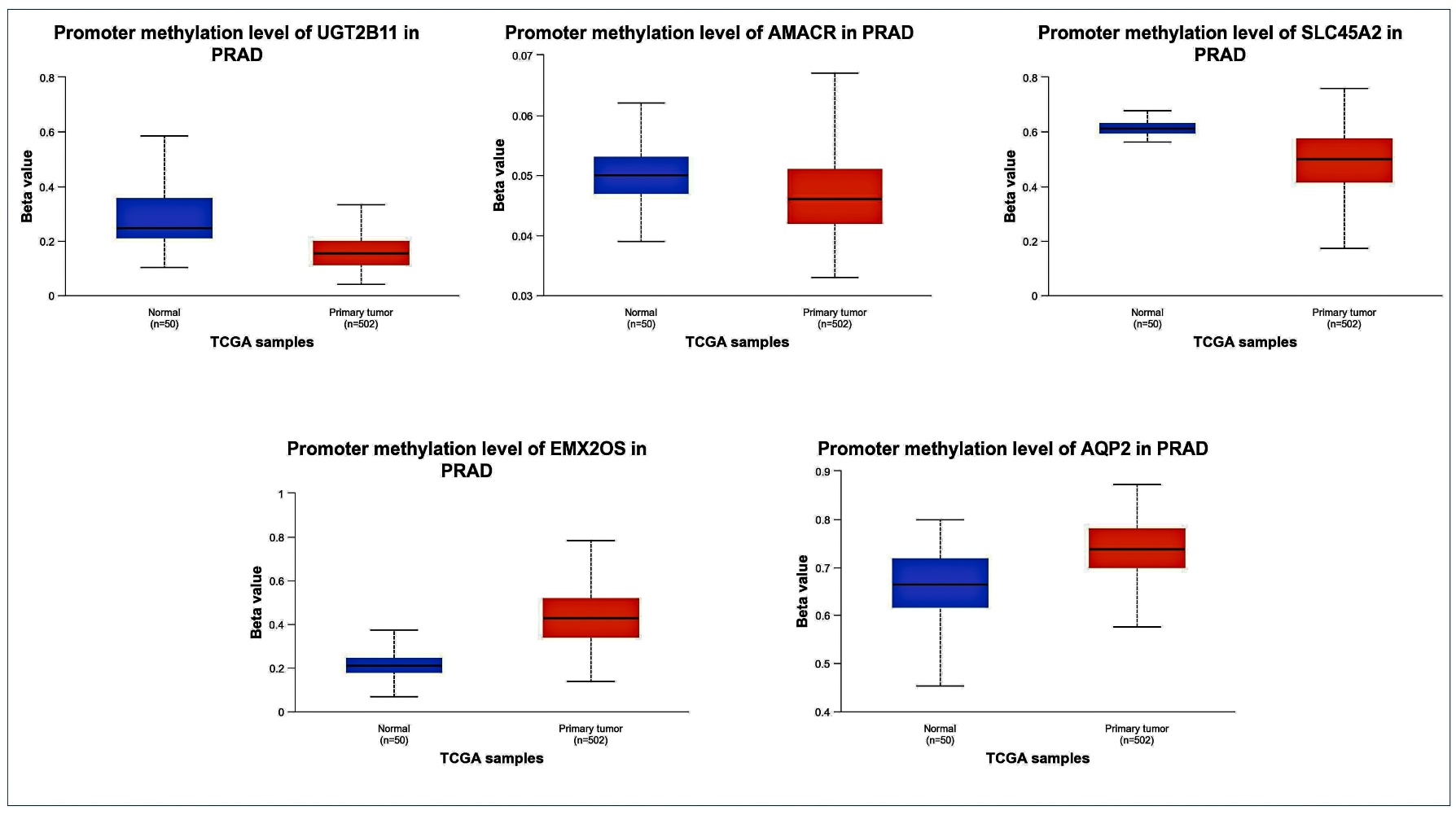

Figure S15. The significant change in promotor DNA methylation levels of UGT2B11 (UGT2B4), AMACR, SLC45A2, EMX2OS, and AQP2 in PRAD.

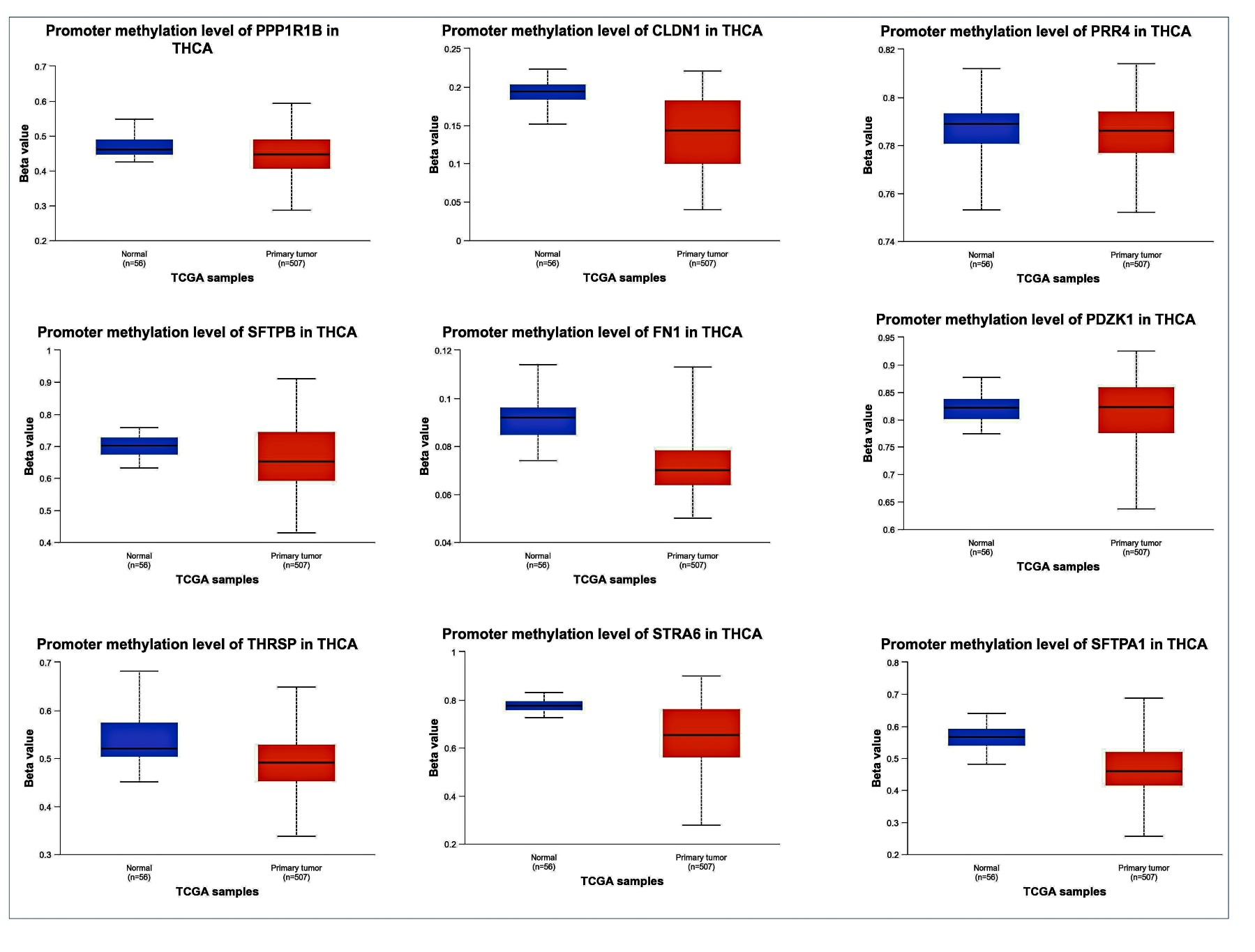
Figure S17. The significant change in promotor DNA methylation levels of SFTPA1, PVRL4, PPP1R1B, STRA6, SFTPB, PDZK1IP1, THRSP, FN1, and CLDN1 in THCA.

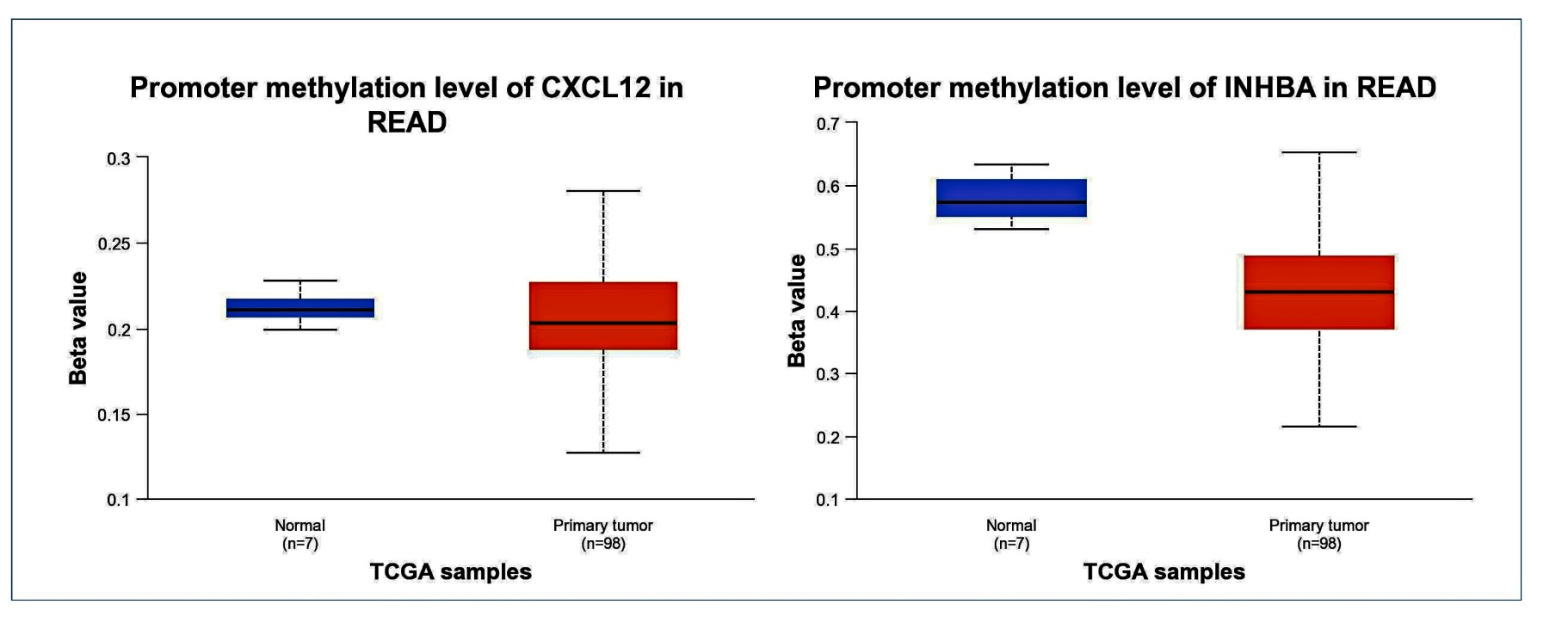

Figure S18. The significant change in promotor DNA methylation levels of CXCL12 and INHBA in READ.
